# Elucidating continental-wide phylogeographic and adaptive processes shaping the genome-wide diversity of North America’s most widely distributed tree

**DOI:** 10.1101/2025.07.08.663217

**Authors:** Roos Goessen, Nathalie Isabel, Christian Wehenkel, Javier Hernández-Velasco, Eduardo Mendoza-Maya, Cuauhtemoc Saenz-Romero, Arnulfo Blanco-Garcia, Jean Bousquet, Ilga M. Porth

**Author notes:** <u>Authors for correspondence:</u> Prof. Ilga M. Porth; Dr. Roos Goessen.

## Abstract

Past population dynamics during the Pleistocene ice age and the Holocene era have profoundly influenced the genetic structure and diversity of species. Environmental heterogeneity has further shaped local and regional adaptive variation. Here, we ask how historical processes have led to the current genetic diversity of a key North American species across its vast natural range and what genomic signatures indicate regional adaptive divergence and local adaptation. We used sequencing data from 1,903 *Populus tremuloides* Michx. (quaking aspen) trees to assess historical population dynamics and identify genotype-environment associations within and among the species’ major genetic lineages. The two northern and western North American aspen lineages exhibited historical population expansion patterns, while the southernmost lineage experienced a historical bottleneck consistent with past glacial oscillations. We found that the earliest split between genetic lineages of *P. tremuloides* occurred in the southern part of its distribution range. We further identified larger blocks of adaptive SNPs within separate genomic sequence regions on chromosomes 2 and 8 that may exhibit suppressed genetic recombination, contributing to the maintenance of regional and local adaptation in the species. Our study provides key insights into the evolutionary processes affecting adaptive genetic variation and phylogeography at a broad continental and regional scale, with implications for predicting species’ responses to future climate change.

## INTRODUCTION

Understanding historical population demographics and movement has been a key theme in species’ evolutionary biology. This knowledge not only sheds light on current genetic population structure but also aids in interpreting the projected population dynamics under anticipated future climates (Jaramillo-Correa *et al*., 2009; Ding *et al*., 2017). Phylogeography, in particular postglacial recolonization routes and vicariance factors affecting geographic structure in North American tree species, have been shown to play a major role in the shaping of a species’ current genetic makeup (Jaramillo-Correa *et al*., 2009). However, questions remain to what extent the local and regional adaptative divergence can be attributed to phylogeography. We investigated the most widespread North American tree species, *Populus tremuloides* Michx., which, given its large continental distribution in nature, may be the key to answering these questions. Indeed, the species ranges across the boreal forests of North America (NA), at higher elevations (>500 m) and in riparian microhabitats in its western range, and in isolated high-elevation habitats (>2000 m) in Mexico and Texas (Little, 1971; Nunneley *et al*., 2014; Goessen *et al.,* 2022). Our previous research revealed four major genetic lineages with strong geographic structure, spanning northeast North America (NE-NA, east of Manitoba and New England), northwest North America (NW-NA, west of Manitoba to Alaska), western US (WU, western US to Baja California), and Mexico (MX, northern and central Mexico) (Goessen *et al*., 2022). Several genetic substructures have also been identified, including a coastal lineage east of the Cascades Mountain range in the Pacific Northwest (Bagley *et al*., 2020; Goessen *et al.,* 2022), and a major north-south subdivision in the NE-NA lineage, suggesting the presence of several glacial refugia in this region during the Last Glacial Maximum (LGM) (Goessen *et al.,* 2022). However, to date, no explicit analyses incorporating genetic information have been conducted to model historical population demography across the range of this keystone species.

Tree species exhibit intraspecific patterns of variation in phenotypic traits, and it has been shown that the underlying genetic variation is largely shaped by local adaptation (Savolainen *et al*., 2007; McKown *et al*., 2014; Porth *et al*., 2015; Depardieu *et al*., 2021; Capblancq *et al*., 2023). Common garden experiments that assess intraspecific variation in traits, including ecophysiology, leaf phenology, and growth-related traits that serve as proxies for fitness, are of great value for assessing patterns of local adaptation (Savolainen *et al*., 2007; McKown *et al*., 2014). However, studies relying on common gardens are expensive and resource-intensive to conduct and often require many years to evaluate traits across the lifespan of trees. Associations between environmental variables and genetic variants*, i.e.* landscape genomics, may offer an alternative approach to assessing certain genetic aspects of local adaptation (Sork *et al*., 2013). The resulting environmental associations can then be used to predict whether natural populations might show genomic maladaptation under future climate conditions and what genetic variations are needed to adapt to such new conditions, as previously shown in *Populus balsamifera* (Fitzpatrick & Keller, 2015), a widespread poplar species found in the boreal forests of North America. However, there is little research on local adaptation at a range-wide scale for species with very large natural distributions such as *P. tremuloides*, given that it would be highly relevant to understand how phylogeography may have contributed to these patterns of divergent adaptive variation. Since *P. tremuloides* can persist on such a large geographic scale with a variety of climatic zones, we expect the presence of a vast number of genetic signatures of adaptation throughout its range. Moreover, our recent study identified around 1,000 *F*_ST_ outlier SNPs with associations to both temperature and precipitation variables within *P. tremuloides*’ four major genetic lineages (Goessen *et al*., 2022). A study on the closely related *P. tremula* also uncovered around 1,000 SNPs associated with environmental variables (Ingvarsson & Bernhardsson, 2019), including a cluster of SNPs located around the *PtFT2* locus, which controls the timing of bud set (Wang *et al*., 2018).

The best-fitting demographic models estimated that *P. tremuloides*, together with its close relative, the Eurasian *P. tremula*, split from a common ancestor around 2.3 million years ago (mya), coinciding with the separation of the Bering land bridge and climate oscillations during the Pleistocene (Wang *et al*., 2016a). The authors studied two populations (Alberta, Canada; Wisconsin, US) and found that *P. tremuloides* has been undergoing population expansion since c.70,000 years ago. It has also been suggested that *P. tremuloides* survived repeated cycles of glaciations in refugia in stable habitats with only minimal displacements along elevational clines or through range shifts (Ding *et al*., 2017; Bagley *et al*., 2020). Strong evidence was based on genetic structure, species distribution models, and ecological niche modelling for refugia in stable habitats that may have persisted at least since the LGM in the western range of the species and in Mexican mountain ranges (Callahan *et al*., 2013; Ding *et al*., 2017; Bagley *et al*., 2020). Separate refugia may also have persisted in the eastern US, from where postglacial expansion may have occurred (Ding *et al*., 2017; Bagley *et al*., 2020; Goessen *et al.,* 2022). Based on common postglacial expansion patterns of several tree species (Jaramillo-Correa *et al*., 2009; Cinget *et al*., 2015; Warren *et al*., 2016), it was assumed that one refugium persisted south of the Great Lakes, more precisely, west of the Appalachian Mountains. Subsequently, postglacial expansion may have occurred from there both northeastward and westward across the North American continent. The other refugia may have persisted east of the Appalachian Mountains, with postglacial expansion toward the north (Jaramillo-Correa *et al*., 2009; Lemieux *et al*., 2011; Cinget *et al*., 2015). Furthermore, the study by Ding *et al*. (2017) found strong evidence for postglacial dispersal from the eastern refugia to the west by evaluating quantitative traits (height, bud break and leaf senescence); in fact, a gradient of decreasing genetic variance in quantitative traits was found from southeast to northwest, reflecting repeated founder events along the historical population migration. Another commonly assumed location for a glacial refugium is the ice-free Beringia land bridge (including Alaska) for several North American species (Edwards *et al*., 2000; Jaramillo-Correa *et al*., 2009; Shafer *et al*., 2010); this has indeed been established for *P. balsamifera* based on palaeobotanical data (Brubaker *et al*., 2005; Breen *et al*., 2012) and has also been suggested for black spruce based on cytoplast and mitochondrial DNA (Gérardi *et al*., 2010). However, paleoclimatic habitat projections for *P. tremuloides* found very little support for the presence of an Alaskan refugium during the LGM (Ding *et al*., 2017), consistent with the lack of geographic structure within this region (Latutrie *et al*., 2016; Goessen *et al*., 2022). There are no palaeobotanical records of *P. tremuloides* in Beringia; pollen from this species is rarely preserved in peats or lake sediments (Brubaker *et al*., 2005); moreover, general pollen identification for poplars beyond the genus level is difficult (Ding *et al*., 2017).

Our previous work aimed to create a global portrait of the genetic makeup of aspen, which was evaluated based on population structure, isolation-by-distance using spatial principal component analysis, population genetic measurements (between-lineage *F*_ST_, heterozygosity, nucleotide diversity), ploidy, clonality, and signatures of selection within genetic lineages (Goessen *et al.,* 2022). The results provided clues for hypotheses to be tested, and the work presented here.

In the present study, which used a larger sample size (1,903 compared to 1,375 aspen individuals in Goessen *et al.,* 2022), we used genome-wide genetic markers to gain insights into the evolutionary history and the shaping of adaptive genetic variation within *P. tremuloides* on a continental scale. First, we sought to reconstruct a population-specific *F*_ST_ phylogeny to better understand population genetic structure in an evolutionary context. With much higher sampling coverage and a broader range than any previous study (Callahan *et al*., 2013; Bagley *et al*., 2020), we then sought to draw statistical phylogeographic conclusions based on models using coalescent theory to expand and refine current theories of postglacial population dynamics and timing since the LGM. We hypothesized that the Mexican and western US *P. tremuloides* lineages may have experienced a stable habitat scenario since the LGM, while we expected expansion in the NE-NA and NW-NA lineages. Moreover, we hypothesized that since the LGM, there has been a recolonization by aspens from multiple refugia in the eastern US, near the Appalachian Mountains and south of the Great Lakes (Jaramillo-Correa *et al*., 2009), towards the western Canada and from a ghost group (Beringia). We also tested for the influence of postglacial migration from the western US lineage or ghost group (presumably from Beringia) with the main migration from the NE-NA lineage, as well as a scenario in which all three groups (ghost population from Beringia, NE-NA, and WUS) come into contact to form the NW-NA lineage. In addition, we wanted to investigate adaptation to local and regional climates for the species’ entire distribution range, using the four major geographic lineages that had previously been delineated at the genetic level (Goessen *et al*., 2022). We asked which environmental factors most strongly drive genome-wide adaptive variation in *P. tremuloides*, and whether these adaptive responses exhibit lineage-specific patterns that reflect their historical biogeographic isolation. We hypothesized that adaptive SNPs associated with climate variables would show differential patterns across the four major genetic lineages (NE-NA, NW-NA, WU, MX), reflecting their historical refugial isolation and subsequent local adaptation trajectories (de Lafontaine *et al*., 2018).

Insights into historical population movements can help explain the current genetic composition of species and provide us with information about how tree populations have responded to past climate oscillations. Findings from locally adaptive genetic variations can also help us predict the future viability of local tree populations by conducting studies that examine genetic incompatibility under changing climate conditions. Moreover, angiosperms are of particular interest for improving the resilience of forests against to increasing fire intensity and frequency, as species such as *P. tremuloides* have been shown to serve as firebreaks and can regenerate quickly after a fire (Shinneman *et al*., 2013).

## MATERIAL AND METHODS

### Study material

*Populus tremuloides* trees can reproduce sexually through seed, and vegetatively through root suckering (clonal reproduction). Moreover, the species can naturally occur as a diploid or triploid. In this study we assessed 1,903 *P. tremuloides* individuals originating from 110 natural populations (**Fig. 1a, b**). Populations ranged up to 420 km and contained 2 to 91 individuals prior to filtering, with the largest pre-filtering population sizes in Mexico, because of their high levels of clonal genotypes (Goessen *et al.,* 2022; Hernández-Velasco *et al*., 2025). Within Mexico, samples were generally taken from clonal stands of 10 individuals within a radius of 200 m (up to a maximum of 4.5 km) (see Hernández-Velasco *et al*., 2025; Goessen *et al.,* 2022), while in other parts of the distribution range, trees were sampled at a distance of at least 500 m. Clonal genotypes were therefore removed using a relatedness filter (see below).

**Figure 1a-c.**
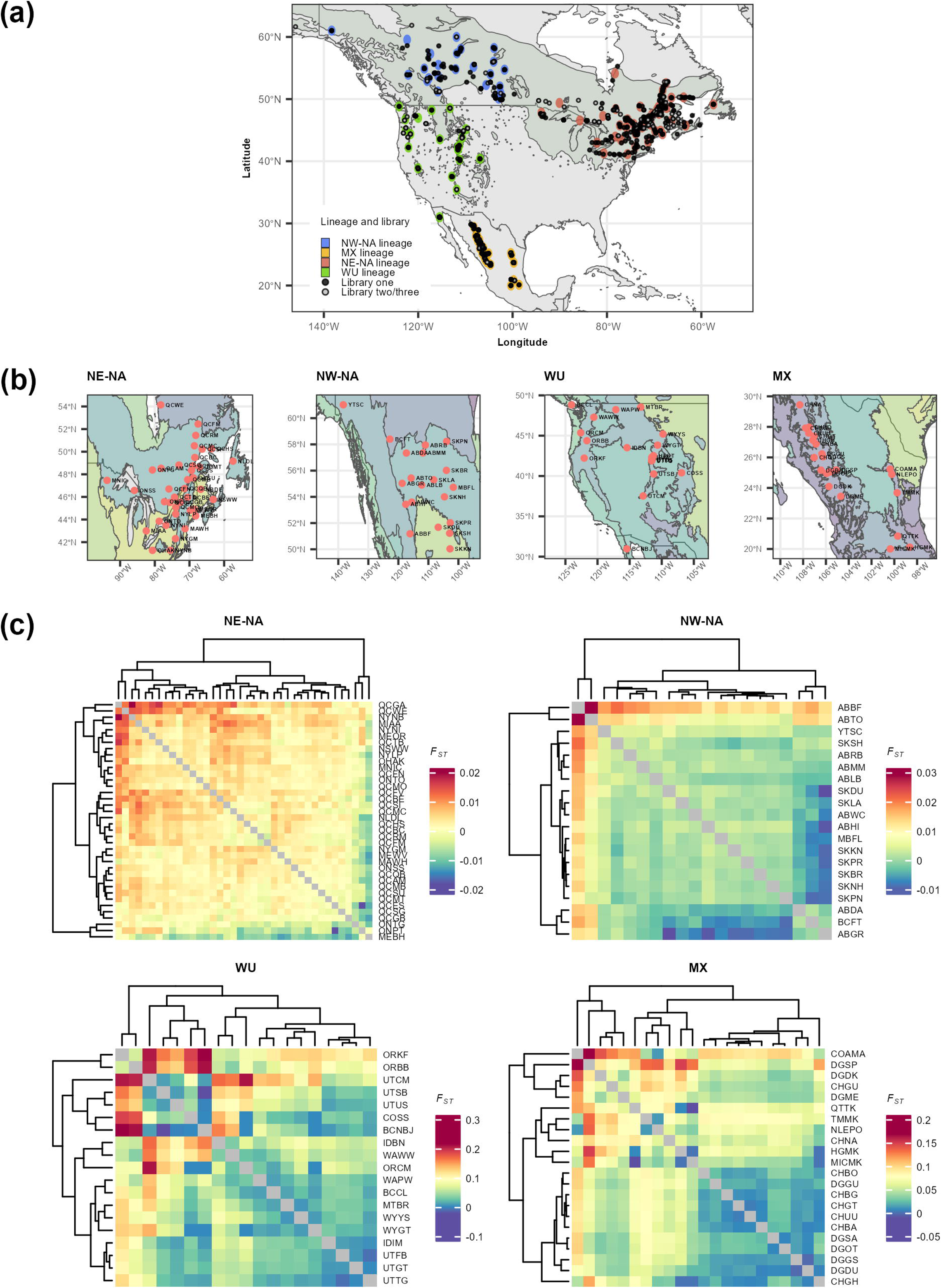
Distribution of all *Populus tremuloides* samples used in this study prior to filtering, genetic lineage assignment and within-lineage *F*_ST_ values. **a.** Distribution of individual samples indicated by library (sequencing library) and assignment of populations to genetic lineages, with genetic lineages MX (Mexico), WU (western US), NW-NA (northwest North America) and NE-NA (northeast North America). **b.** Map displaying the location and name of populations within each genetic lineage. **c.** *F*_ST_ heatmap within each genetic lineage on a population-basis.

### Genomic data preparation

*P. tremuloides* (**Fig. 1a, b**) and *P. grandidentata* (81 individuals used to check for spontaneous hybrids or misclassified individuals) were gathered, DNA isolated, and sequenced as described (Goessen *et al*., 2022; see **Dataset S1**). One additional DNA library with 550 individuals was sequenced (using a single lane on Illumina Novaseq S4) that included samples with a DNA concentration of 10ng/ul (n=189) as well as samples with a DNA concentration of 3ng/ul (n=361) for triple digest genotyping-by-sequencing (GbS) library preparation using restriction enzymes PstI/NsiI/MspI (3D-GbS). SNPs were called as described in Goessen *et al*. (2022), with modifications including the usage of: 1) 110 reclassified populations for SNP calling (populations step in STACKS) as determined by Goessen *et al*., 2022; 2) our new *P. tremuloides* reference genome to improve SNP calling and obtain chromosomal positions of SNPs. Related to point 2), a SNP calling comparison between the new and the old reference (plantgenie.org, v1.1) was performed (**Methods S1**, **Table S1**).

For ploidy determination, we first filtered the raw SNPs with VCFtools (v0.1.16) (Danecek *et al*., 2011) under very loose settings to obtain a large number of SNPs (--remove-indels, maf 0.05, --max-missing 0.5, --min-meanDP 12, --max-meanDP 100). Ploidy was determined with FastPloidy with cut-off 0.9 (Goessen *et al*., 2022). In total, 320 samples were removed due to too low heterozygous SNP count (below 1000) for clear ploidy determination. The FastPloidy ratio is based on the allelic ratio (number of reads mapped to heterozygous SNPs), with triploids having a ratio above 1, and diploids having a ratio below 1 (see Goessen *et al.,* 2022 for further details). Samples that were in a range between 0.9-1.1 were removed due to ambiguity in ploidy (n=20) and samples above the 1.1 cut-off were called as triploid. In total, 170 triploids and 1,474 diploids were identified (see **Dataset S1**).

Triploids, ploidy ambiguous and identified *P. grandidentata* samples (as described in Goessen *et al.,* 2022) were eliminated from the raw dataset. Duplicate samples from the second library were also removed. Only SNPs on the 20 major scaffolds were kept (remaining contigs and haplotigs were removed). The data was then filtered using a filtering pipeline available at https://github.com/enormandeau/stacks_workflow with the settings described in **Table S2.** We decided to only work with the first library for demographic analyses because we observed a difference in observed heterozygosity between duplicate samples present in both libraries. For GEA (genotype–environment association) analyses, we considered both libraries but with more stringent missingness and MAS (minimum number of samples with rare alleles) settings to reduce the difference in heterozygosity.

### Pairwise *F*_ST_ calculations and phylogenetic tree construction

The pairwise population differentiation estimate *F*_ST_ (Weir & Cockerham, 1984) was determined using the Hierfstat pairwise.WCfst() function in R, over the entire species range as well as based on the species’ previously inferred (Goessen *et al.,* 2022) genetic lineages. The pairwise *F*_ST_ heatmaps were visualized with R package ggplot2. For the entire species’ range, a UPGMA tree with *F*_ST_ values including bootstrapping (10,000 simulations) was constructed using the R package ape and visualized with R package ggtree. The pairwise *F*_ST_ matrix was implemented in SplitsTree4 to create a phylogenetic neighbour joining tree. In addition, we determined the maximum likelihood phylogeny using the GTRGAMMA model and 1,000 bootstrap replicates in RAxML v8.2.9 (Stamatakis, 2014) on the entire species’ range. RAxML was run with 6,000 random SNPs from Set_01 and converted to phylip format using PGDspider. The resulting tree was visualized using R package ggtree with “equal angle” layout.

The phylogeny and *F*_ST_ analyses were based on the genetic lineages as determined by Goessen *et al*. (2022). Based on preliminary results, we regrouped the BCCL population (from Vancouver Island) into the western US (WU) lineage. In addition, we removed the NBDE population (6 individuals from New Brunswick) from our dataset because this population showed extreme divergence in the phylogeny.

### Assessment of demographic history from genetic polymorphisms

#### Fastsimcoal2 to infer the demographic history of each genetic lineage

The folded site frequency spectra (SFS) were generated with the EasySFS.py script (https://github.com/isaacovercast/easySFS, version 2022-04-26) and downsampled to 156, 84, 212, 391 for lineages MX, WU, NW-NA and NE-NA, respectively. Demographic models were tested using fastsimcoal version 2.7 (Excoffier *et al*., 2021) with 50 optimization cycles (-L) and 200,000 simulations (-n) per likelihood estimate. SFS entries with less than 10 SNPs were pooled to limit overfitting (-C). Parameter estimates were selected based on the estimated likelihood that had the smallest difference with the maximum observed likelihood out of 100 runs in fastsimcoal2. The Akaike information criterium (AIC) was used to compare the likelihoods of the best run and identify the most likely model. We used a per-site mutation rate of 2.5x10^-9^ (Tuskan *et al*., 2006; Wang *et al*., 2016a) and a generation time of 15 years (Wang *et al*., 2016a).

We tested, for each of the four genetic lineages, a stable, a bottleneck, an expansion, and a contraction model (**Fig. S1**) to assess demographics since the LGM. The models were run with a minimum effective population size of 10 000 in the .est file. The “stable” scenario had a fixed timespan that calculated if the effective population size remained stable over the last 2,000 generations (*i.e.* 30,000 years), to test for a stable effective population size during and after the LGM. Moreover, we tested for the inclusion of the “bounded” keyword to specify if the upper time range is bounded (or not) during parameter estimation (see **Methods S2**).

#### Random Forests (RF) based Approximate Bayesian Computation (ABC) to deduce the demographic history among the genetic lineages

We used DIY-ABC-RF to derive the choice of the demographic model and the inference of parameters (Collin *et al*., 2021) (**Methods S3**). The training set was generated with 4,213 SNPs after selecting those with minor allele frequency (MAF) of 0.05 from a set of 6,000 randomly picked SNPs. We first ran the program for model selection with only 10 simple scenarios (Run1), to determine the order of splits between the major genetic lineages. Hereafter, we ran the program a second time for model selection using 13 scenarios (Run2), including the best scenarios of Run1 along more complex scenarios that follow the same split order of lineages. Lastly, we ran the program another time with only the most likely scenario from Run2 for parameter estimation (Run3).

For model selection, we performed 2,000 simulations per scenario in the training set (*i.e.* 20,000 simulations for Run1 and 26,000 simulations for Run2) and used 1,500 trees for the random forest analysis. The scenario with the highest classification votes in a forest of 1,500 trees represents the most likely scenario among the set of all tested scenarios. For Run3, we performed 40,000 simulations and again used 1,500 trees for parameter estimation of the most likely scenario.

The scenarios in Run1 (for shorthand, R1S1-R1S10) are visualized in **Fig. S2**, and do not display any admixture events or influence of ghost scenarios. We chose to always split the MX from the WUS lineage, as these lineages are genetically and geographically closest (Goessen *et al.,* 2022). The scenarios in Run2 (R2S1-R2S13) include the most likely simple scenarios without admixture events, as well as scenarios with admixture and additional ghost scenarios (**Fig. S3**). In detail, for Run2, R2S1 and R2S2 describe the two most likely scenarios of Run1 (see the Results section for interpretation), *i.e.* a split of the MX and WUS lineages at T3, followed by a split by either WUS and NWNA (*i.e.* R2S1 = R1S1) or WUS and NENA (*i.e.* R2S2 = R1S2) at T2, and a split of NENA and NWNA at T1. R2S3 describes an admixture event between the WU and NE-NA lineages that formed the NW-NA lineage. R2S4 describes the influence of a ghost lineage G1 formed from the WU lineage (potentially the Beringia ghost population) after splitting from the NE-NA lineage; G1 then recombined with the NE-NA lineage to form the NW-NA lineage. R2S5 describes the same as R2S4; however, here, the ghost lineage was formed before the potential diversion between the WU and NE-NA lineages. R2S6 describes the influence of a ghost lineage G2 that split from the NE-NA lineage before recombining with the WU lineage to form the NW-NA lineage. G2 represents a ghost lineage that was potentially located south of the Great Lakes during the LGM, while other lineages of the original NE-NA lineage were assumed east of the Appalachian range. R2S7 and R2S8 describe the influence of both the ghost lineage that diverged from the WU lineage (G1), as well as the influence of the NE-NA and WU lineages to form the NW-NA lineage. For R2S7, G1 diverged at T2a, while for R2S8, G1 diverged at T2. R2S9 and R2S10 describe the divergence of a ghost lineage (G2) and following its recombination with the NE-NA lineage, as well as admixture between WU and NE-NA, further followed by an admixture event with G1 to finally form the NW-NA lineage. R2S9 describes the split of G1 and WU at T2b (after the split from the NE-NA lineage), while R2S10 describes the split of G1 and WU at T2 (before the split from the NE-NA lineage). R2S11 and R2S12 describe the split of a ghost lineage from WU followed by an admixture event between WU and the ghost lineage to form respectively the NW-NA or NE-NA lineage. R2S13 describes a split of a ghost lineage from the NW-NA lineage that subsequently had an admixture event with the NW-NA lineage to create the NE-NA lineage.

As historical parameter priors, we used the following: C (ancestral effective population size) [40 000, 70 000], NE-NA [40 000, 500 000], NW-NA [40 000, 500 000], WU [20 000, 500 000], MX [10 000, 500 000], G1 [10 000, 500 000], G2 [10 000, 500 000], T1 [500, 2000], T2 [2000, 50 000], T3 [3000, 160 000], T1a [300, 2000], T1b [300, 2000], T2a [2000, 50 000], T2b [2000, 50 000], ra [0.01, 0.99], rb [0.01, 0.99], rc [0.01, 0.99]. The upper boundaries for population size were based on results by Wang *et al*. (2016a), who found a current effective population size of 309 500 in aspen populations from Manitoba, while the ancestral population size was found to be 56 235 (2.33 mya). We decided to assign lower prior boundaries for the WU and MX lineages than for the NW-NA and NE-NA lineages, as we expected potential lower effective population sizes since these populations are more isolated. The upper boundary for T1 (in number of generations) was determined based on the timing of the LGM, while the upper boundary for T3 was based on the split of *P. tremuloides* and *P. tremula* around 2.3 mya. Moreover, the timepoints were forced to follow the order depicted in **Figs. S2-3**, *e.g*., T1 cannot be earlier than T2 etc.

We evaluated whether scenarios and associated priors were compatible with the observed dataset using a visual assessment of the projection of the simulated datasets vs. observed dataset on the first linear discriminant axes (LDA) axes (Pudlo *et al*., 2016). The observed dataset should be within the cloud of simulated datasets (Pudlo *et al*., 2016). Moreover, to evaluate the quality of the DIY-ABC-RF predictions for scenario choice, the global prior error rates are reported. This includes a contingency table of predicted vs. true scenarios as well as the mean classification error rate over all scenarios. Lastly, we report the posterior error of the selected scenario by RF. The tested scenarios using DIY-ABC-RF were compatible with the dataset, as shown by the visual assessment of the simulated datasets vs. the observed datasets on the first LDA (linear discriminant analysis) axes, *i.e*. the observed dataset was within the cloud of the simulated datasets (**Figs. S4-5**). The DIY-ABC-RF headerRF.txt files are uploaded as supplementary data.

### Genotype–environment association analyses

First, environmental data per individual was extracted from the ClimateNA database, version 7.32 (Wang *et al*., 2016b). This database downscales climate normal data (800x800m) to scale-free point locations (Wang *et al*., 2016b). We extracted the 1961-1990 climate normals for 24 available annual climate variables (see **Dataset S2**, in spreadsheet format Excel, for all climate variables per individual); abbreviations of variables can be found in **Table S3**. We then calculated the principal components (PCs) over all environmental variables for our GEA analyses in order to obtain insights into the overall patterns of adaptive variation, and also performed the GEA analyses with all individual climate variables in order to gain more background information. When running our GEA analyses, we assessed one variable at a time, and thus variables were not all grouped together in a single run. We decided to consider PC3, given that its percentage of explained variation was close to that of PC2 for range-wide variation. For consistency, we therefore considered this PC for analyses conducted within genetic lineages. All variables were scaled by subtracting the mean and dividing through the standard deviation across all samples. We additionally conducted the GEA analysis using a less stringent MAS filter and employing non-collinear variables for the principal component (PC) calculation (see **Methods S4**). This was done to ensure that the original MAS filtering criteria did not inadvertently exclude important signals of adaptation.

We used both Latent factor mixed models 2.0 (LFMM2) (Caye *et al*., 2019) from the LEA R package (Gain & François, 2021) and the Bayesian program Bayenv2 (Günther & Coop, 2013) to perform GEA analyses. The GbS library number (1,2,3) was considered in the LFMM2 analysis and not scaled. The library number was not considered in the Bayenv2 analysis as it runs on population means. As LFMM2 does not accept missing data, we performed imputation using a script (developed by M. Lamothe at NRCan) that uses a machine learning approach within the R package XGBoost (Chen & Guestrin, 2016) with the following parameters: booster = "dart", eta = 0.2, gamma = 0, subsample = 0.8, objective = "multi:softmax", nrounds = 200. We chose to run LFMM2 with 5 latent factors that represent the four major lineages and the substructure in the western US lineage, based on the variance explained by each principal component implemented in the LFMM2 package. Resulting *P* values were adjusted using the Benjamini-Hochberg correction to reduce false positives. SNPs with an adjusted *P* value below 0.05 were considered to be significantly associated with the tested environmental variable. Moreover, we ran LFMM2 separately within each of the four major genetic lineages (using one latent factor). Because only bi-allelic (no mono-allelic) SNPs are accepted by the program, we extracted 12669, 12647, 13097 and 10247 SNPs, respectively, for the NE-NA, NW-NA, WU and MX lineages.

We ran Bayenv2 using the entire sample size, as well as separately within each genetic lineage. We averaged the climate variables per population to run this program, and the climate PCs were calculated over these means. We first estimated a covariance matrix using Bayenv2 with 100,000 Markov Chain Monte Carlo (MCMC) runs. We then ran Bayenv2 with 100,000 MCMC runs using the -c option that includes Pearson and Spearman correlations of allele frequencies with the environmental data. Resulting SNPs that displayed a Bayes factor above 30, indicating very strong evidence for the alternative hypothesis, were considered as having a significant association with the tested variable. Moreover, to reduce the selection of false positives, we considered only SNPs that were part of the top 5% of SNPs with significant environmental correlations based on both Spearman and Pearson coefficients. Further description of the differences between the two approaches is discussed in **Methods S5** (with the related dataset).

Blastx was employed using the *P. tremuloides* CDS sequences to obtain identifiers for *A. thaliana* (highest hit below an E-value of 1e^-10^). Moreover, GO terms and GOSLIM terms were extracted for *A. thaliana* identifiers using the ATH_GO_GOSLIM.txt file available at the *Arabidopsis* Information Resource (TAIR, https://www.arabidopsis.org) website (last updated on 2023-03-01). GO enrichment analysis was performed on SNPs associated with PC1, PC2 and PC3, separately for Bayenv2 and LFMM2, using AgriGo (Tian *et al*., 2017). Gene descriptions for *A. thaliana* identifiers were extracted from the TAIR database as well as the UniProt database (https://www.uniprot.org). We used BlastKoala to assess KEGG (Kyoto Encyclopedia of Genes and Genomes) ontology terms for genes identified *via* their associations with PC1, PC2 and PC3, and based on blasting the protein sequences against a nonredundant set of KEGG GENES (Kanehisa *et al*., 2016). Using the BlastKoala assignment, we assessed if the identified SNPs were in or near genes that are present in the same biological pathways. Additionally, we assessed the overlap with Bayescan outlier genes from Goessen *et al*. (2022). We ran Interproscan v5.52-86.0 (Jones *et al*., 2014) on all genes that were close or within SNPs associated with climate PC1, PC2 and PC3 identified within each genetic lineage. Only one PFAM and SUPERFAMILY per gene was retained based on the lowest E-value. Resulting annotations were categorized per genetic lineage, and those occurring more than once per genetic lineage were visualized using the treemap R package. Moreover, genes that were close or within SNPs associated with PC1, PC2 and PC3 identified within each genetic lineage were further categorized using literature search, and separately for biological process and molecular function.

Finally, we used R package gaston (Dandine-Roulland & Perdry, 2017) to calculate and visualize linkage disequilibrium (LD) using the r^2^ measure for SNPs identified for PC1, PC2 and PC3 based and individual environmental variables over the entire sample with either LFMM2 or Bayenv2.

### Environmental SNPs and sequence inversion data

We used Oxford Nanopore Technologies (ONT, PromethION) data of four *P. tremuloides* individuals (Goessen *et al.,* in prep; Porth *et al*., 2024a), representing the major genetic lineages in aspen (Goessen *et al*., 2022), to determine whether SNPs with environmental associations match the location of the DNA sequence inversions. The ONT individuals are named FP2.5-13 (origin: Mexico, Durango, latitude = 23.5, longitude = -104.7; the same individual was also used to construct the reference genome), FBR-3672-A (origin: US, Utah, latitude = 42.0, longitude = -111.6), PNL-9 (origin: Canada, Saskatchewan, latitude = 58.2, longitude = -103.9), and ESSI-001-O-A (origin: Canada, Quebec, latitude = 48.3, longitude = -69.4).

The ONT data from each sample were aligned against the *P. tremuloides* reference genome using NGMLR (v0.2.7) (Sedlazeck *et al*., 2018) with default parameters. The alignment BAM files were analyzed by Sniffle2 (v2.0.2) (Smolka *et al*., 2024) to identify structural variations with precise breakpoints and minimum number of 20 reads supporting the structural variation. Inversions and SNPs with environmental associations were visualised using the R package Circlize.

## RESULTS

### *F*_ST_ phylogeny and heatmaps for aspen

Populations within NE-NA and NW-NA lineages displayed relatively low *F*_ST_ values (respectively up to 0.02 and 0.03), while populations within the WU and MX lineages showed higher *F*_ST_ values (respectively up to 0.22 or 0.17) (**Fig. 1c**). The overall heatmap with *F*_ST_ values between all populations had *F*_ST_ values ranging between -0.02 to 0.36 (with DGSP (Durango) vs. ORCM (Oregon) populations showing the highest value) (**Fig. S6a**). The maximum-likelihood phylogenetic tree shows that individuals are grouping into their respective lineages (**Fig. S6b**).

### Demographic history within genetic lineages in aspen

Across all simulations run in fastsimcoal2 at the genetic lineage level, expansion was the most likely scenario for NE-NA, NW-NA and WU lineages, while bottleneck the most likely scenario for the MX lineage (**Table 1, Table S4, Fig. 2A).** NE-NA, NW-NA and WU experienced expansion since, respectively, 5383, 5387 and 1883 generations, translating to 80,745, 80,805 and 28,245 years ago (**Table 1**). Effective population sizes were found to have increased from 10,307 to 17,820 for NE-NA, from 10,116 to 17,665 for NW-NA and from 10,524 to 15,913 for WU. The MX lineage experienced a strong bottleneck 1019 generations ago (*i.e.* 15,285 years ago), thereby declined in effective population size from 190,913 to 1323 and subsequently increased again (to 85,004 in the calculation).

**Figure 2a-b.**
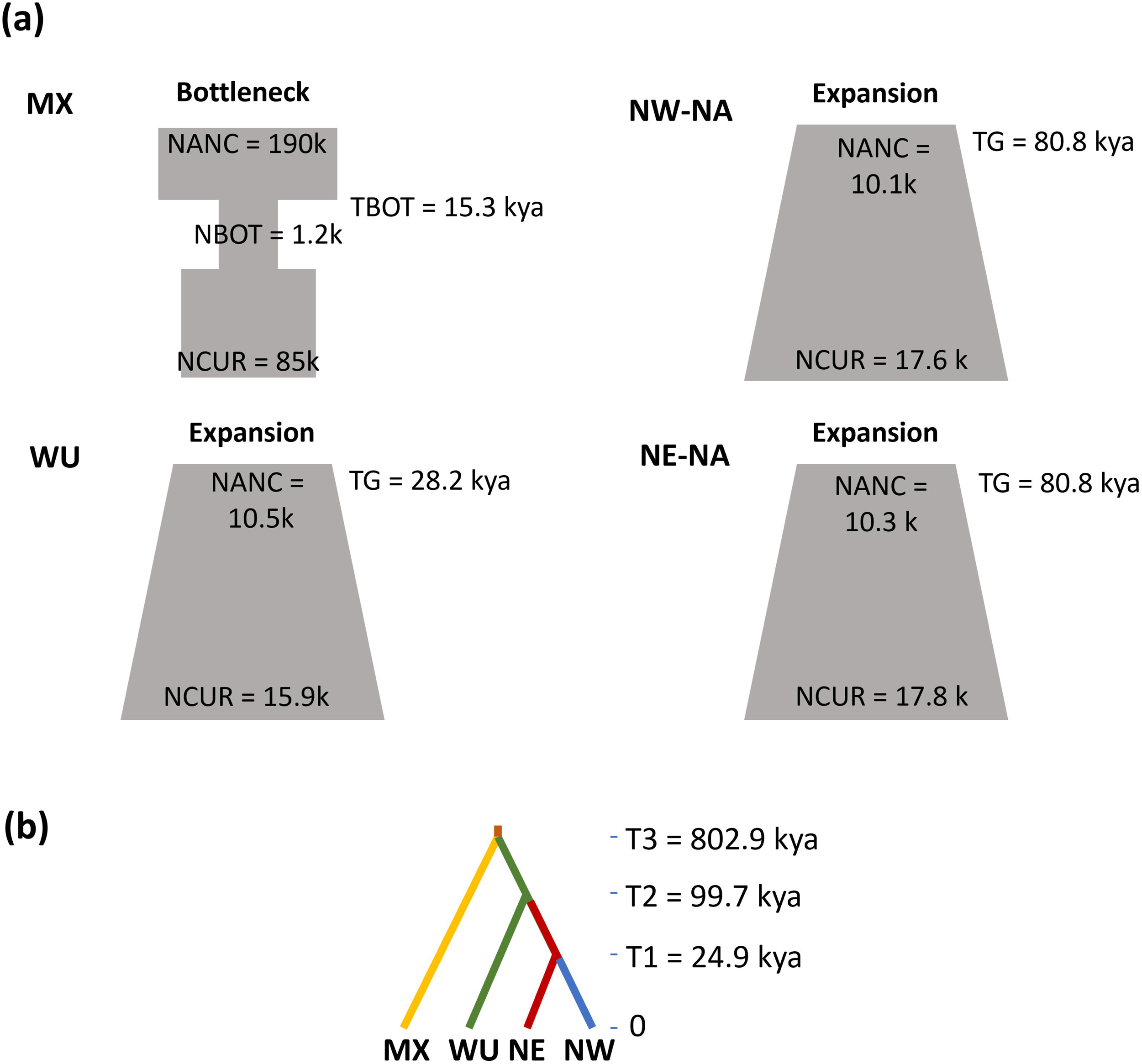
Most likely scenarios within and between lineages. **a.** The most likely within-genetic lineage scenario following fastsimcoal2 analyses (**Fig. S1**), with the following genetic lineages: MX (Mexico), WU (western US), NW-NA (northwest North America) and NE-NA (northeast North America). NANC indicates ancient population size, NCUR indicates current population size, TBOT indicates start of bottleneck in generations, NBOT indicates population size during bottleneck, TG indicates the number of generations since population expansion or contraction. **b.** The most likely between-genetic lineage scenario following DIY-ABC-RF analyses, *i.e.* scenario 2 in both, Run1 (**Fig. S2**) and Run2 (**Fig. S3**).

**Table 1.** Statistical results for best model runs of demographic scenarios tested within a single lineage with fastsimcoal2, without using “bounded” keyword. Bold rows indicate the most likely demographic scenario for said lineage based on lowest AIC (Akaike Information Criteria) values. DeltaL indicates the difference between the maximum observed likelihood (under a perfect fit of the expected to the observed SFS) and maximum estimated likelihood under the model parameters. Lineages included northeast North America (NE-NA), northwest North America (NW-NA), western US (WU) and Mexico (MX). Np indicates number of parameters included in the model. NCUR indicates the current effective population size, TG indicates the number of generations since start of expansion/contraction, NANC indicates ancient effective population size, GROWTHRT indicates the growth rate, NBOT indicates effective population size during the bottleneck, TBOT indicates the number of generations at which the bottleneck started. TG_15 indicates the generation time of TG in number of years (TG * 15 years). TBOT _15 indicates the generation time of TBOT in number of years (TBOT * 15 years).

| Lineage | Scenario | Np | deltaL | AIC | NCUR | NANC | TG | TG_15 | GROWTHRT | NBOT | TBOT | TBOT_15 |
| --- | --- | --- | --- | --- | --- | --- | --- | --- | --- | --- | --- | --- |
| NE-NA | Stable | 1 | 3888.3 | 213619.0 | 108427.0 |  |  |  |  |  |  |  |
|  | Bottleneck | 4 | 997.6 | 200307.0 | 363175.0 | 1021759.0 |  |  |  | 1378.0 | 11.0 | 165.0 |
|  | Contraction | 4 | 988.9 | 200266.8 | 14120.0 | 368005.0 | 1470.0 | 22050.0 | 0.00222 |  |  |  |
|  | **Expansion** | **4** | **823.9** | **199506.8** | **17820.0** | **10307.0** | **5383.0** | **80745.0** | **-0.00010** |  |  |  |
| NW-NA | Stable | 1 | 2735.2 | 184628.0 | 164495.0 |  |  |  |  |  |  |  |
|  | Bottleneck | 4 | 717.9 | 175337.9 | 450851.0 | 921699.0 |  |  |  | 1543.0 | 12.0 | 180.0 |
|  | Contraction | 4 | 711.9 | 175310.0 | 14762.0 | 412391.0 | 1455.0 | 21825.0 | 0.00229 |  |  |  |
|  | **Expansion** | **4** | **551.2** | **174570.3** | **17665.0** | **10116.0** | **5387.0** | **80805.0** | **-0.00010** |  |  |  |
| WU | Stable | 1 | 393.7 | 127767.4 | 222579.0 |  |  |  |  |  |  |  |
|  | Bottleneck | 4 | 67.0 | 126262.8 | 11740.0 | 698802.0 |  |  |  | 3933.0 | 24.0 | 360.0 |
|  | Contraction | 4 | 65.9 | 126257.7 | 21219.0 | 425965.0 | 1234.0 | 18510.0 | 0.00243 |  |  |  |
|  | **Expansion** | **4** | **24.6** | **126067.7** | **15913.0** | **10524.0** | **1883.0** | **28245.0** | **-0.00022** |  |  |  |
| MX | Stable | 1 | 490.9 | 101408.9 | 90270.0 |  |  |  |  |  |  |  |
|  | **Bottleneck** | **4** | **9.2** | **99190.7** | **85004.0** | **190913.0** |  |  |  | **1232.0** | **1019.0** | **15285.0** |
|  | Contraction | 4 | 92.3 | 99573.4 | 22337.0 | 36052.0 | 529.0 | 7935.0 | 0.00091 |  |  |  |
|  | Expansion | 4 | 90.9 | 99566.9 | 22964.0 | 21112.0 | 266.0 | 3990.0 | -0.00032 |  |  |  |

### Demographic history between genetic lineages in aspen

Using the RF scenario choice for Run1 (in the DIY-ABC-RF approach), which included only simple scenarios aimed at identifying the initial split among lineages, scenario 2 (R1S2) received the highest support, gathering 709 classification votes out of a total of 1,500 trees, corresponding to a posterior probability of 0.696 (posterior error of 1-0.696=0.307) (**Table S5**). R1S1 followed with 654 votes, and R1S5 was the third most likely scenario but with only 30 votes (**Table S5**). R1S2 describes a divergence history in which MX and WU lineages split first from a common ancestor at time T3, followed by the separation of the WU and NE-NA lineages at T2, and finally the split between the NE-NA and NW-NA lineages at T1.

Using the RF scenario choice on Run2, that included the top two most likely scenarios of Run1 as well as more complex scenarios with ghost clusters and admixture events, the highest number of classification votes went to R2S2 (the same scenario as in Run1, thus R1S2) with a total of 370 classification votes, under a posterior probability of 0.625 (posterior error of 1-0.625=0.375). The second most likely choice was R2S13 with 359 classification votes (**Table S6**). Across both, Run1 and Run2, the majority of misclassifications occurred between scenario 2 and scenario 1, and *vice versa* (**Table S7-S8**). The misclassification error per scenario ranged from 0.003 to 0.37 and from 0.21 to 0.61 (**Table S7-S8**), for respectively Run1 and Run2.

Following the parameter estimation for scenario 2 (R2S2) (**Table 2**, **Fig. 2B**), the MX and WU lineages separated 53,530 generations ago, *i.e.* 802,950 years ago (under 15-year generation time, posterior mean), with 0.05-0.95 quantiles ranging between 453,670 and 1,154,589 years ago. The WU and NE-NA lineages separated 99,660 years ago (0.05-0.95 quantiles ranging between 50,008 and 164,655 years ago), and the NE-NA and NW-NA lineages separated 24,885 years ago (0.05-0.95 quantiles ranging between 16,950 and 29,745 years ago). The ancestral effective population size at 802,950 years ago was 55,802. Parameter estimations for the current effective population sizes display a posterior mean of 233K, 56K, 375K and 278K for the lineages MX, NE-NA, NW-NA and WU, respectively (**Table 2**). Quantiles (0.05-0.95) for effective population size show ranges of 107-462K, 41-78K, 207-491K, and 128-472K, respectively for lineages MX, NE-NA, NW-NA and WU.

**Table 2.** Parameter prediction for the most likely tested scenario, *i.e.* scenario 2, in DIY-ABC-RF. C indicates ancestral effective population size. MX, NE-NA, NW-NA and WU indicate the effective population size respectively for the genetic lineages Mexico (MX), northeast North America (NE-NA), northwest North America (NW-NA) and western US (WU). T1 indicates time point 1, T2 indicates time point 2 and T3 indicates time point 3; all in generations ago. Year indicates the number of generations multiplied by the generation time of 15 years.

| Variable | Posterior mean | Year (x15) | Median | Year (x15) | Quantile_0.05 | Quantile_0.95 | Variance |
| --- | --- | --- | --- | --- | --- | --- | --- |
| C | 55802.1 |  | 56735.1 |  | 41481.9 | 68461 | 6.65E+07 |
| MX | 233477 |  | 199319 |  | 107327 | 462680 | 4.09E+09 |
| NE-NA | 56066.2 |  | 52134.2 |  | 41176.6 | 77934.1 | 1.06E+08 |
| NW-NA | 375536 |  | 388669 |  | 207621 | 491810 | 7.20E+09 |
| WUS | 278960 |  | 245346 |  | 128059 | 472165 | 4.76E+09 |
| T1 | 1659.81 | 24885 | 1728 | 25920 | 1130 | 1983 | 45229.8 |
| T2 | 6644.37 | 99660 | 6211.75 | 93165 | 3333.86 | 10977 | 4.65E+06 |
| T3 | 53530.9 | 802950 | 52244.6 | 783660 | 30244.7 | 76972.6 | 9.61E+07 |

### Rangewide-scale environmental associations in aspen

The highest number of SNPs identified with the Bayenv2 method (**Dataset S3**, **Fig. 3**) encompassed 72 SNPs for the MCMT climate variable, while the LFMM2 method (**Dataset S4**, **Fig. 3**) identified 178 SNPs associated with variable bFFP. No SNPs were associated with library number in LFMM2. Among all unique SNPs identified by either method, the highest numbers of SNPs were identified on scaffolds 3, 13 and 15 of the aspen genome (**Table S9**), respectively representing chromosomes 2, 8 and 16 (following the *P. alba* genome synteny).

**Figure 3.**
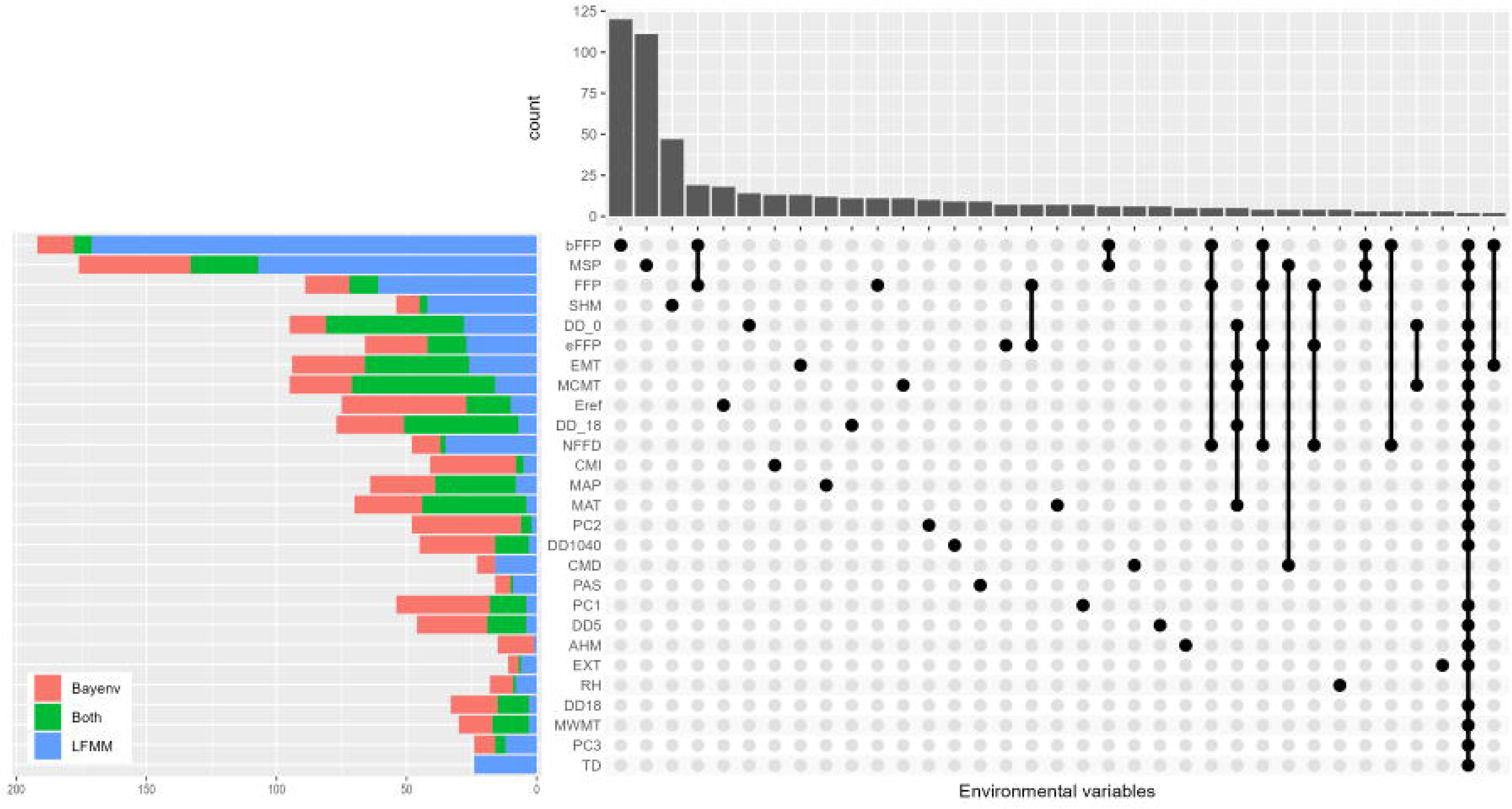
Depiction of SNP counts overlap between the different individually tested environmental variables and climate PC1, PC2 and PC3 (identified over the entire sample). Upset plot with lower columns indicating if and how the SNP sets associated with the different variables intersect, while the upper bar plots depict the SNP counts per set (= set sizes). A maximum of 35 columns (x-axis) in the upset plot is shown. The number of SNPs identified per GEA analysis method and variable (left plot) is indicated in red for Bayenv2, blue for LFMM2 and green for both methods.

PC1, PC2 and PC3 explained, respectively, 58.5%, 15.5% and 12.1% of the variation in climatic variables (**Fig. S7-8**). Across the entire sample distribution, the strongest correlation for PC1 was with MAT (r=0.98), for PC2 with MAP (r=-0.85), and for PC3 with MWMT (r=-0.9) (**Table 3**). Using Bayenv2, 50, 46, and 12 SNPs were identified for PC1, PC2 and PC3, respectively (**Fig. 3**, **Dataset S4**). Using LFMM2, 18, 6, and 16 SNPs were identified for PC1, PC2 and PC3, respectively (**Fig. 3**, **Dataset S4**). The overlap between the two methods resulted in 14, 4 and 4 SNPs, respectively, for PC1, PC2 and PC3. For Bayenv2, 26 SNPs were identified for at least two PCs, and 4 SNPs for all three. For LFMM2, 15 SNPs were identified for both PC1 and PC3. For PC1, most identified SNPs were located on scaffolds 3 and 13, as shown by the peaks of the adjusted *P* values as well as the Bayes factor for LFMM2 and Bayenv2, respectively (**Fig. 4**). Also on aspen genome scaffold 3, clear peaks of high significance for PC2 and PC3 remain visible using both Bayenv2 and LFMM2 (**Fig. 4**). Out of all the 84 SNPs identified with both methods for PC1, PC2 and PC3, 19 were missense variants, further 19 were synonymous variants, 8 were intron variants, and the remainders were either downstream, upstream, or intergenic variants. Out of all the SNPs identified by Bayenv2 and LFMM2 for PC1, PC2 or PC3, 34 SNPs were also identified through *F*_ST_ outlier analysis (using the Bayescan approach) and reported previously by Goessen *et al*. (2022) (see **Dataset S5, Table 4**). The GEA analyses conducted with a lower MAS filter and non-collinear variables for PC calculation yielded results consistent with the original analysis, showing similar peaks of identified SNPs for PC1, PC2, and PC3 on scaffolds 3 and 13 (see **Results S1**, **Figs. S9-17**, **Table S10**).

**Figure 4a-f.**
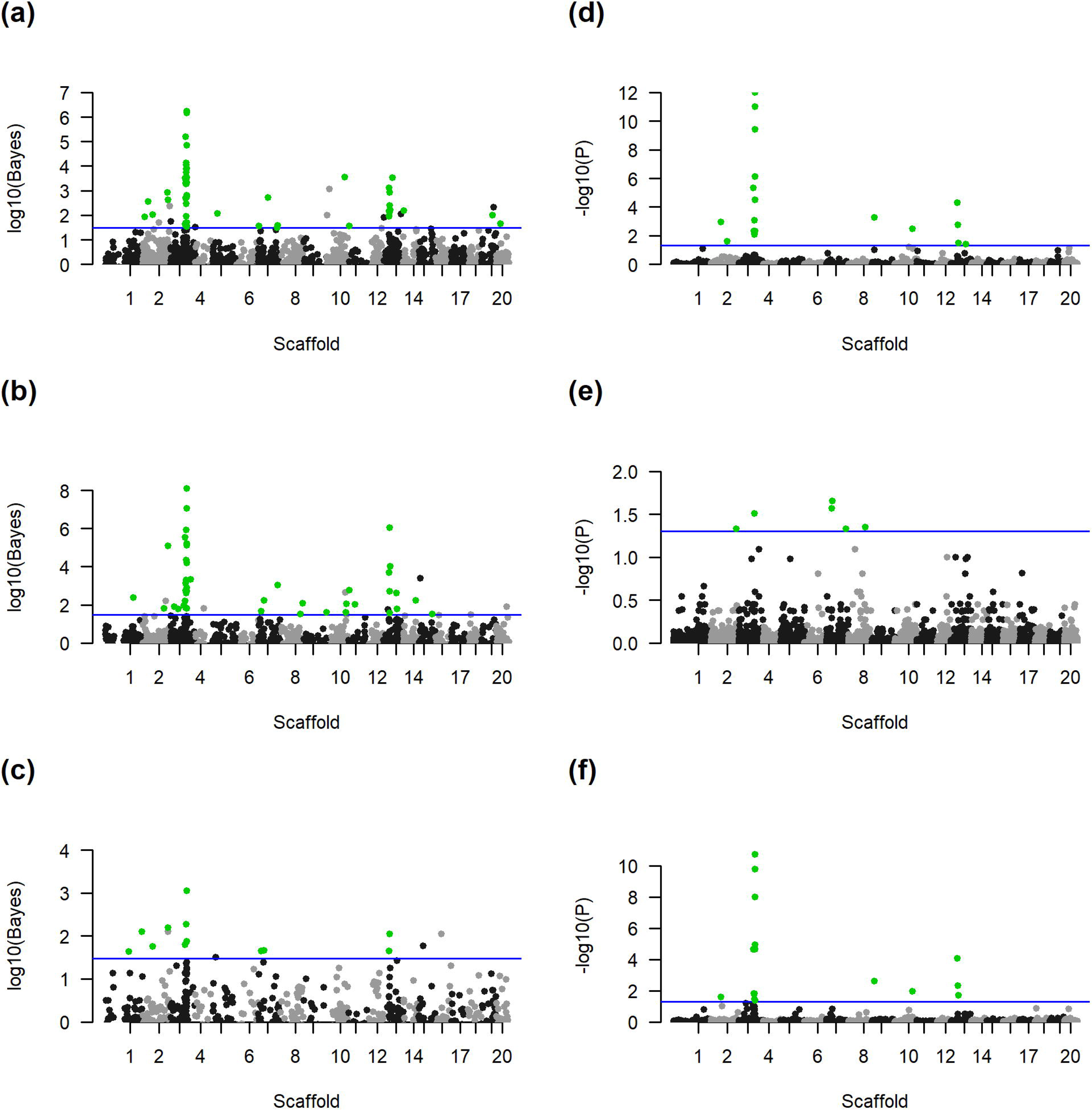
Manhattan plots of results for Bayenv2 (Bayes factor, figs. a-c) and LFMM2 (adjusted *P* value (*P*), figs. d-f) for PC1 (figs. a, d), PC2 (figs. b, e) and PC3 (figs. c, f) for all samples along the 20 major scaffolds. The blue line for Bayenv2 plots indicates the cut-off Bayes factor of 30. The blue line for LFMM2 indicates the adjusted *P* value cut-off of 0.05. SNPs identified by Bayenv2 that are above the blue cut-off line, but not colored green (Figs. **a-c**) did not pass the criteria for overlap with the top 5% for Pearson and Spearman correlations.

**Table 3.**
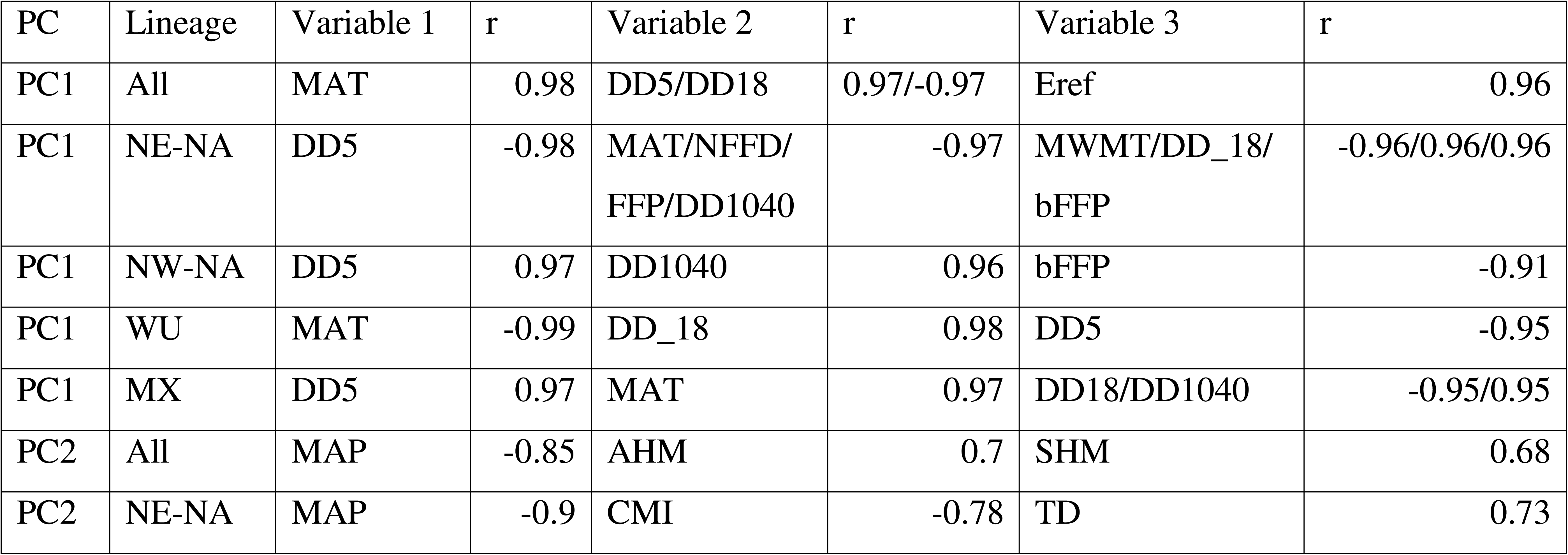
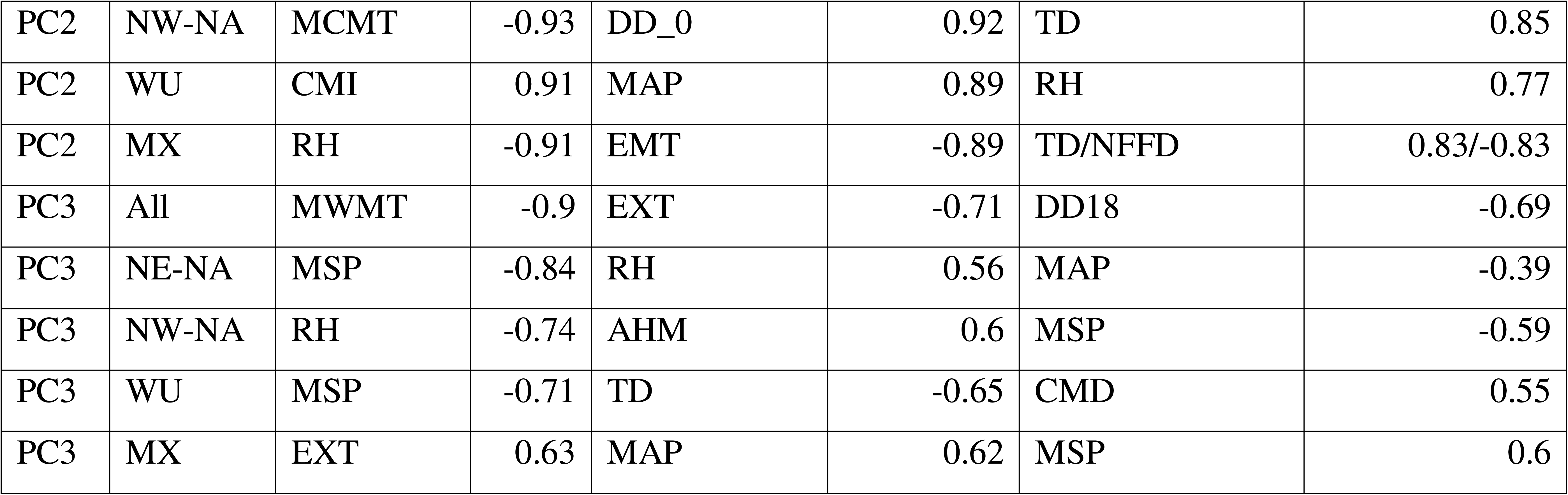
Top three strongest correlations (r) of PC1, PC2 and PC3 with climatic variables. Tested climatic parameters of geographic origins (Wang *et al*., 2016b) include the following: mean annual temperature (MAT, °C), mean warmest month temperature (MWMT, °C), mean coldest month temperature (MCMT, °C), temperature difference between MWMT and MCMT (TD, °C), mean annual precipitation (MAP, mm), May to September precipitation (MSP, mm), annual heat-moisture index (AHM, (MAT+10)/(MAP/1000)), summer heat-moisture index (SHM, (MWMT/(MSP/1000)), degree-days below 0°C (DD_0), degree-days above 5°C (DD5), degree-days below 18°C (DD_18), degree-days above 18°C (DD18), number of frost-free days (NFFD), frost-free period (FFP), the day of the year on which FFP begins (bFFP), the day of the year on which FFP ends (eFFP), precipitation as snow (PAS, mm), extreme minimum temperature over 30 years (EMT), extreme maximum temperature over 30 years (EXT), Hargreaves reference evaporation (Eref, mm), Hargreaves climatic moisture deficit (CMD, mm), mean annual relative humidity (RH, %), Hogg’s climate moisture index (CMI, mm), degree-days above 10°C and below 40°C (DD1040). PC indicates principal component. Correlations were calculated over the entire sample, as well as within each genetic lineage, *i.e.* northeast North America (NE-NA), northwest North America (NW-NA), Western US (WU) and Mexico (MX).

| PC | Lineage | Variable 1 | r | Variable 2 | r | Variable 3 | r |
| --- | --- | --- | --- | --- | --- | --- | --- |
| PC1 | All | MAT | 0.98 | DD5/DD18 | 0.97/-0.97 | Eref | 0.96 |
| PC1 | NE-NA | DD5 | -0.98 | MAT/NFFD/<br>FFP/DD1040 | -0.97 | MWMT/DD_18/<br>bFFP | -0.96/0.96/0.96 |
| PC1 | NW-NA | DD5 | 0.97 | DD1040 | 0.96 | bFFP | -0.91 |
| PC1 | WU | MAT | -0.99 | DD_18 | 0.98 | DD5 | -0.95 |
| PC1 | MX | DD5 | 0.97 | MAT | 0.97 | DD18/DD1040 | -0.95/0.95 |
| PC2 | All | MAP | -0.85 | AHM | 0.7 | SHM | 0.68 |
| PC2 | NE-NA | MAP | -0.9 | CMI | -0.78 | TD | 0.73 |
| PC2 | NW-NA | MCMT | -0.93 | DD_0 | 0.92 | TD | 0.85 |
| PC2 | WU | CMI | 0.91 | MAP | 0.89 | RH | 0.77 |
| PC2 | MX | RH | -0.91 | EMT | -0.89 | TD/NFFD | 0.83/-0.83 |
| PC3 | All | MWMT | -0.9 | EXT | -0.71 | DD18 | -0.69 |
| PC3 | NE-NA | MSP | -0.84 | RH | 0.56 | MAP | -0.39 |
| PC3 | NW-NA | RH | -0.74 | AHM | 0.6 | MSP | -0.59 |
| PC3 | WU | MSP | -0.71 | TD | -0.65 | CMD | 0.55 |
| PC3 | MX | EXT | 0.63 | MAP | 0.62 | MSP | 0.6 |

**Table 4.** Annotation of genes on scaffold 3 (chromosome 2) that encompass or are near adaptive SNPs identified with both Bayenv2 and LFMM2 over the entire sample for climate PC1, PC2 and PC3, as well as showing an overlap with genes identified by *F*_ST_ outlier analysis in Goessen *et al*., 2022. PTS: *Populus tremuloides* gene name; ATG: *Arabidopsis thaliana* gene name identified with blastx.

| PTS | ATG | Name ATG | Annotation |
| --- | --- | --- | --- |
| *Pt25377* | *AT1G28200* | *FIP1* | Seed dormancy, germination, response to ABA (Li *et al.*, 2023). |
| *Pt25404* | *AT4G01720* | *WRKY47* | Transcription factor involved in response to water deprivation (Raineri *et al.*, 2015). |
| *Pt25412/Pt25413* | *AT3G62150* | *ABCB21* | Involved in auxin transport (Kamimoto *et al.*, 2012). |
| *Pt25468* | *AT2G47260* | *ATWRKY23* | Transcription factor involved in auxin transport (Prát *et al.*, 2018). |
| *Pt25470* | *AT4G01970* | *RS4* | Involved in raffinose synthase, response to oxidative stress (Nishizawa *et al.*, 2008; Gangl *et al.*, 2015). |
| *Pt25479* | *AT4G02010* | n.a. | Unknown. |

Wide-ranging LD between SNPs identified on scaffold 3 within genomic region 14,611,739–16,388,003 was especially apparent for samples of the NE-NA genetic lineage (**Fig. 5**), the MX lineage and the overall sample (**Fig. S18**), while LD was less strong when calculated within WUS or NW-NA lineages (**Fig. S18**). We highlight the localized distribution of three missense SNPs on this genomic region, namely, positions 15468642, 15562784 and 16268712, corresponding to genes *Pt25404* (homolog of *AtWRKY47*), *Pt25412* (homolog of *AtABCB21*) and *Pt25470* (homolog of *AtRS4*), respectively. All three SNPs showed a shift in their allelic states between eastern North America and the rest of the distribution range (**Fig. 6**). For allele frequency differences among aspen lineages involving another SNP of interest within this region (*LACS1* gene) see Goessen *et al*. (2022).

**Figure 5.**
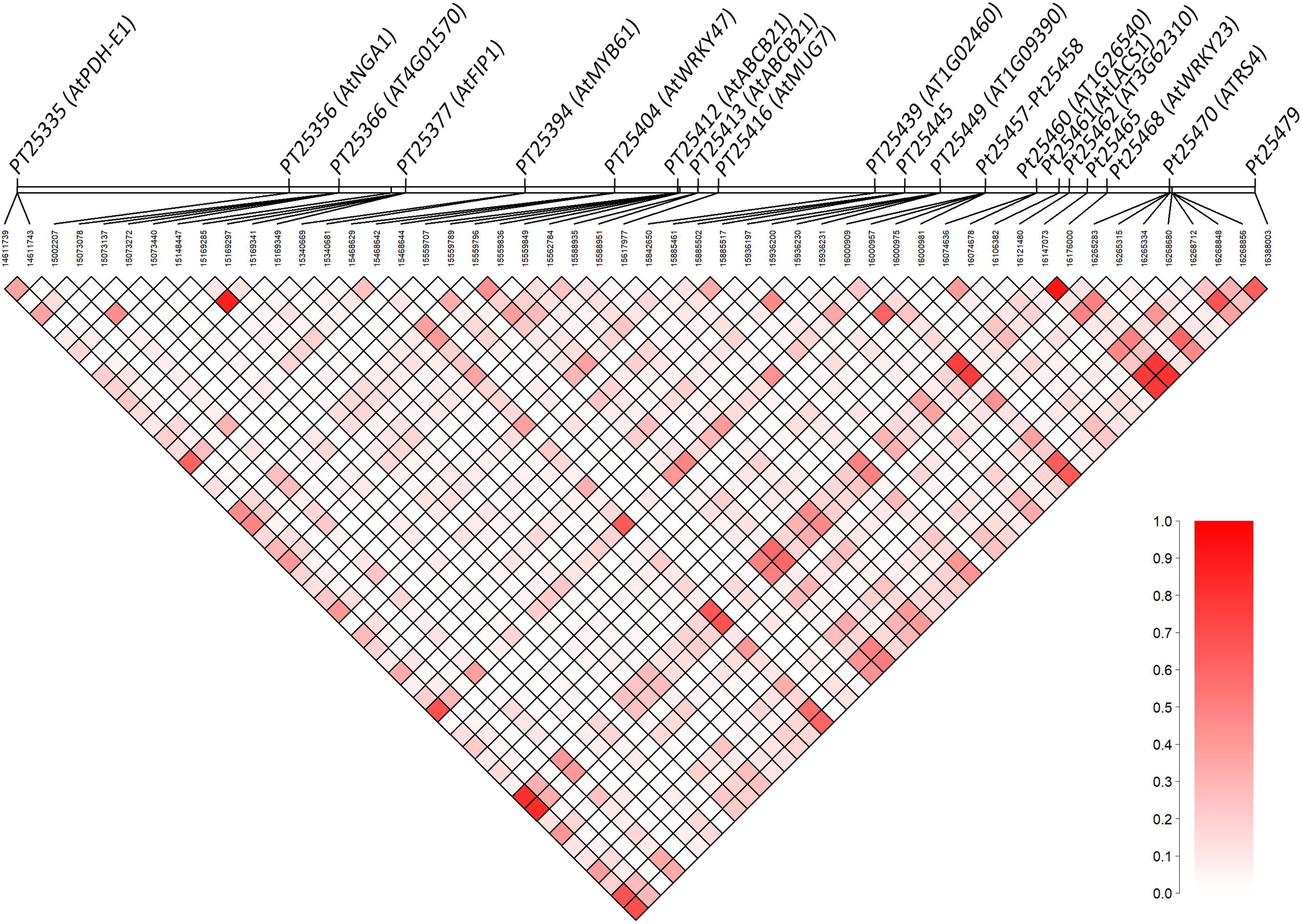
Linkage-disequilibrium (LD, r^2^) plots of SNPs identified in GEA analyses for climate PC1, PC2 and PC3 using Bayenv2 and LFMM2 on a range-wide scale. The plot visualizes the genomic region 14611739 – 16388003 on scaffold 3 (representing chromosome 2 from alignment against *Populus alba*). The plot visualized the LD within the genetic lineage northeastern North America (NE-NA). LD-plots for the northwestern North America (NW-NA), western US (WUS) and Mexico (MX) genetic lineages are visualized in **Fig. S18**. With a color scheme ranging from r^2^=0 in white towards r^2^=1 in red*. Populus tremuloides* gene names are indicated for SNPs near (<5kb) or directly within these genes, with corresponding *Arabidopsis thaliana* gene names based on blastx in brackets. Preliminary results of long-read sequencing four distant individuals (Porth *et al*., 2024a) show a major inversion spanning 9.5 Mbp starting at base pair 11,177,991 (heterozygous) at scaffold 3 in an individual from the NE-NA lineage (compared to the MX individual); moreover, individuals from the MX (heterozygous) and WU (homozygous) lineages show an inversion of only 2.8 Mbp size starting at bp 12,892,408 (Goessen *et al*., unpublished results).

**Figure 6.**
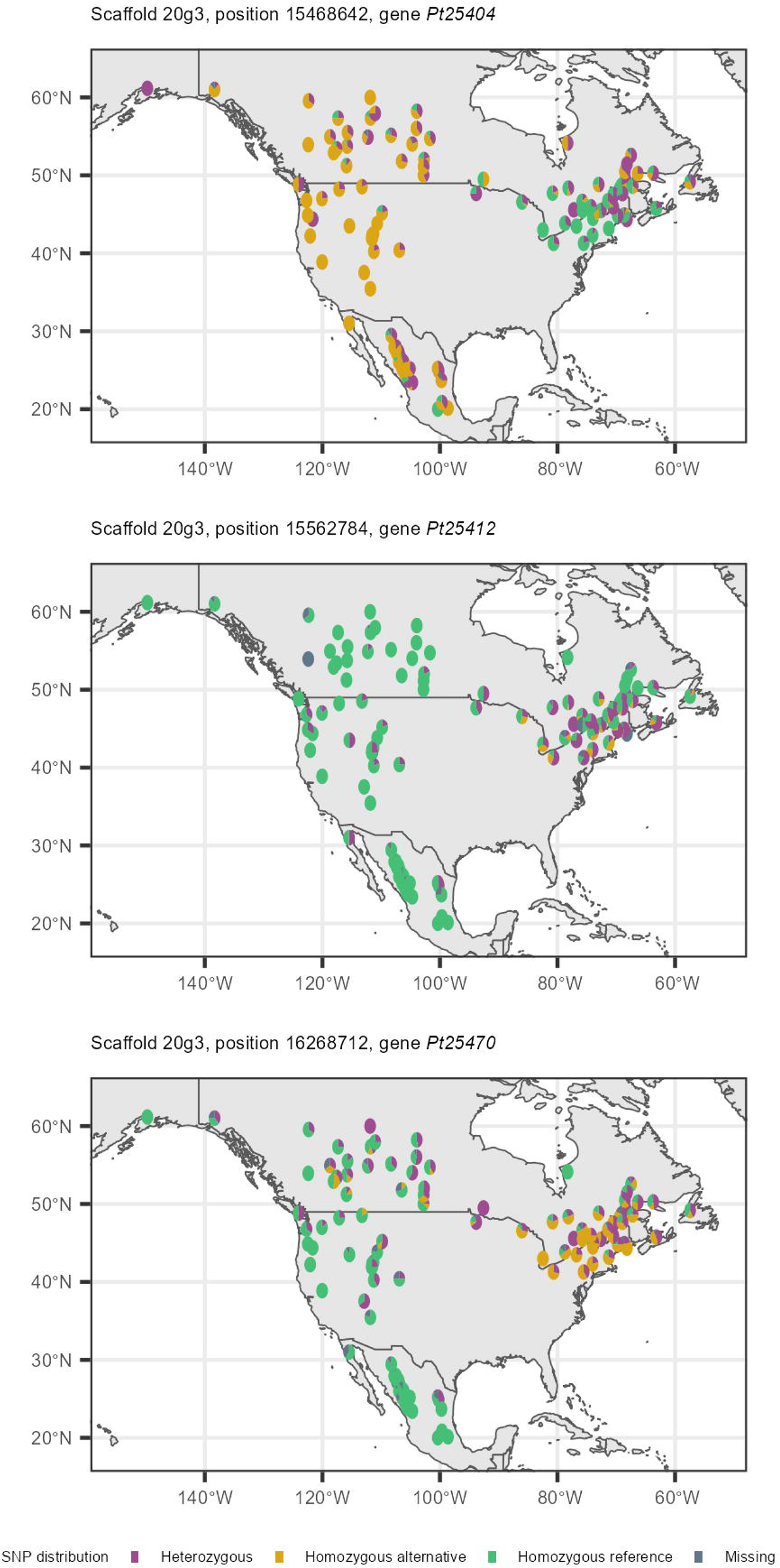
The visual validation of three selected environment-associated SNPs in *Populus tremuloides*. We show the distribution of three missense SNPs on scaffold 3 (chromosome 2) corresponding to genes *Pt25404*, *Pt25412* and *Pt25470*. These genes are respectively homologs of *Arabidopsis thaliana* genes *AtWRKY47*, *AtABCB21* and *AtRS4*.

Inversions overlapping the genomic region on scaffold 3 (positions 14,611,739–16,388,003) were identified in several individuals sequenced with ONT methodology. Specifically, the sequenced ESSI-001-O-A individual (representative of the NE-NA lineage) exhibited a heterozygous inversion spanning genome sequence positions 11,177,991 to 20,743,620; the PNL-9 individual (NW-NA lineage) had a heterozygous inversion from 4,353,721 to 12,879,523; individual FBR-3672-A (WU lineage) showed a homozygous inversion between 12,892,408 and 15,724,881; and finally, FP2.5-13 (MX lineage) carried a heterozygous inversion ranging from 12,892,408 to 15,724,886 (**Dataset S6**, **Fig. 7**).

**Figure 7.**
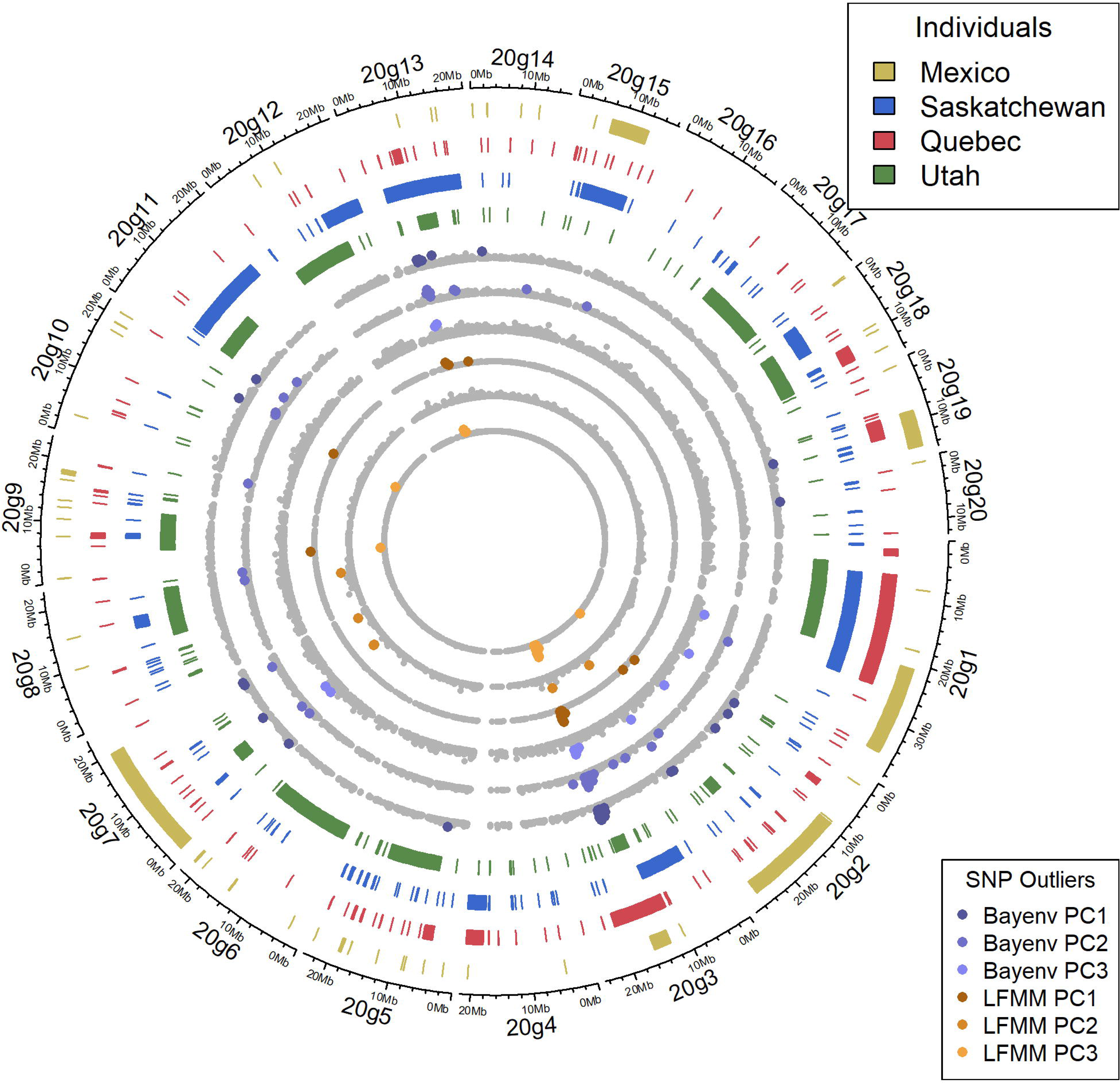
Localization of the inversions identified in four *Populus tremuloides* individuals and the corresponding positions of the SNPs associated with climate PC1, PC2 and PC3 identified over the entire sample. We show that most SNPs with significant associations are located on scaffold 3, in a region that displays inversions among the tested individuals. Displayed SNPs correspond to those visualized in the Manhattan plots of Fig. 4.

### Environmental associations *within* the four genetic lineages of aspen

Using Bayenv2, within the NE-NA lineage, we identified three SNPs for PC1, one for PC2 and again one SNP for PC3 (**Table S11, Dataset S7**). Within the NW-NA lineage we only identified one SNP for PC1. For the WU lineage, two SNPs were identified for PC1, one for PC2 and two for PC3. Within the MX lineage, six SNPs were identified for PC1, one for PC2 and eight for PC3.

Using LFMM2, we identified 25 SNPs for PC1 and two SNPs for PC2 within the NE-NA lineage (**Dataset S7**). In total, 72 SNPs were identified for PC1 in the NW-NA lineage, 36 for PC2 and five for PC3. We did not identify any significant SNPs for the WU lineage related to any of the three PCs. For the MX lineage, we identified two SNPs for PC1 and 50 SNPs for PC3.

The overlap of SNPs between the two methods was very low, with only two overlapping SNPs for PC1 for the NE-NA lineage, and seven SNPs for PC3 for the MX lineage. Interestingly within the NW-NA lineage, 24 SNPs were also associated with library number (LFMM2), of which nine SNPs were overlapping with those associated with climate PC1 and PC2 in that same lineage. However, none of the other lineages showed significant SNPs with library number. Due to the unexpectedly high number of SNPs identified for NW-NA, of which many were also associated with library number, we decided to run the program with only samples of the first GbS library using LFMM2. This resulted in substantially less SNPs identified for the NW-NA lineage, *i.e*. 17 for PC1, 3 for PC2 and 4 for PC3 (**Table S11, Dataset S7**). No overlaps were found among the four lineages in terms of SNPs identified within lineages.

### Annotations for adaptive SNPs in aspen

No significant GO-term enrichments were identified among the SNPs associated through LFMM2 or Bayenv2 for PC1, PC2 or PC3, neither over the entire sample nor within the four genetic lineages. Using BlastKoala, we found that the identified SNPs were not located in or near genes that would suggest the same biological pathways. For further assessment, we explored the functional characterization of *A. thaliana* homologs in aspen. Within the genetic lineages, the number of genes associated with development and or environmental response were 14 out of 24 (58.3%), 7 out of 22 (31.8%), 2 out of 5 (40%) and 21 out of 53 (39.6%), respectively, for the NE-NA, NW-NA, WU and MX lineages (**Fig. S19, Dataset S8**). Additionally, we identified the following gene number for transcription factors within genetic lineages: respectively 4 out of 24 genes (16.7%), 1 out of 22 genes (4.5%), none out of 5 genes (0%), and 5 out of 53 genes (9.4%) for NE-NA, NW-NA, WU and MX lineages (**Fig. S19**). Also, no prominent overrepresentation in the frequency of PFAMs or SUPERFAMILYs among the identified genes within the genetic lineages or over the entire sample was found (**Fig. S20**).

## DISCUSSION

This study provides important insights into the phylogeographic processes and adaptive mechanisms in the context of a keystone species, *P. tremuloides*, widely distributed at the continental scale of North America: (1) First, distinct historical population dynamics were observed within lineages, attributed to the different vicariance and demographic factors in each region during the glacial cycles and following the Holocene. Specifically, the two northern lineages displayed historical expansion patterns that may have followed the penultimate glacial retreat (Wang *et al*., 2016a), while the Mexico lineage displayed a distinct bottleneck pattern conforming with the LGM and possible expansion following the onset of global warming after the LGM. (2) Second, divergence of major lineages in the southern part of North America may have followed the start of extreme glaciation cycles after more systematic global cooling during the first half of the Pleistocene (Snyder, 2016). Based on between-lineage phylogeographic analyses (our DIY-ABC-RF analysis) we identified that the oldest split occurred between the western US and Mexico genetic lineages, potentially indicating that the species’ ancestral populations are genetically closest to individuals from the southern range. (3) Third, local adaptation is likely facilitated by genomic islands of divergence. Identified blocks of adaptive SNPs on aspen chromosome 2 (scaffold 3) and on aspen chromosome 8 (scaffold 13) may be the result of genomic rearrangements bringing together co-adapted loci (Yeaman, 2013); indeed, a large inversion (of several Mbp) spanning these regions in the aspen genomes was identified among four genetically distant individuals using Oxford Nanopore Technologies (ONT) long-read sequencing against the ONT reference assembly used in the current paper.

The close similarity in the identified parameter settings between the NE-NA and NW-NA lineages is in line with the low *F*_ST_ values and close phylogeny between populations of these lineages, and in agreement with the hypothesis of a more recent shared origin and shared postglacial expansion patterns from eastern refugia after the last glacial maximum (Ding *et al*., 2017; Bagley *et al*., 2020; Goessen *et al.,* 2022). While we hypothesized an expansion pattern for the NE-NA and NW-NA lineages starting around 15 kyrs ago representing the retracting of ice sheets since the LGM, the genomic signatures resulting from the penultimate glaciation appeared to be stronger than those following the expansion and retraction of icesheets in relation to the LGM (Wang *et al*., 2016a). In line with our findings, a study on seven major European forest tree species, including *Populus nigra* (section *Aigeiros*), found that effective population size has often been maintained despite range contractions caused by glacial cycles (Milesi *et al.,* 2024); here, authors suggested that the genetic diversity of large tree populations can be quite resilient to sudden environmental change.

Distinct patterns of regional adaptive evolution in aspen were observed within its two northern genetic lineages in comparison to its western US and Mexico lineages. Within the NE-NA and NW-NA lineages, we identified most significant SNPs for PC1, which showed the highest correlation with temperature variables, thus temperature might be a stronger driver of local adaptation in these trees, due to their necessary adaptation for transition from temperate to boreal forests. Similarly, a study in white spruce sampled within northeastern NA, similar to the geographical range of the NE-NA lineage range in aspen, found that most significant SNPs were rather associated with mean annual temperature than mean annual precipitation (Hornoy *et al*., 2015). Moreover, adaptive genetic variation for temperature may be involved in optimizing bud phenology timing (Rohde *et al*., 2011; Olson *et al*., 2012). Compared to other genetic lineages, the NE-NA lineage had relatively the most SNPs in or near genes related to environmental response and development as well as transcription factors, in line with the observed high climatic variability. Previous research identified only very few *F*_ST_ outlier SNPs within the NW-NA lineage in comparison to the other assessed genetic lineages of aspen, attributable to the low environmental variability within this range (Goessen *et al*., 2022); indeed, a relative low number of genes associated with environmental response or development was identified in this lineage compared to the others.

Against our expectations, the WU lineage exhibited a demographic expansion that would not align temporally with the retreat of the LGM ice sheets, which began around 19,000 years ago. In contrast, the inferred timing of expansion for this lineage occurred earlier, around 28,000 years ago, during the onset of glacial conditions rather than following ice retreat. One possible explanation is that the high rate of clonal reproduction in the WU lineage (Mock *et al*., 2008), and similarly in the MX lineage (Goessen *et al*., 2022), could extend generation times, thereby complicating the temporal identification of demographic inferences. While we expected a stable habitat scenario for the WU lineage (Ding *et al*., 2017; Bagley *et al*., 2020), patterns of retraction and expansion along elevational shifts could still be expected following climate oscillations (Callahan *et al*., 2013; Ding *et al*., 2017; Bagley *et al*., 2020). Moreover, the lack of identified SNPs for the WU lineage was unexpected. Potentially, the first three PCs did not capture the most important climatic gradients for local adaptation within this lineage, compared to individual climatic variables, for which many SNPs were found to be associated (*e.g*., DD18, FFP, see supplemental results) within this lineage.

The MX lineage experienced a relatively recent bottleneck event coinciding with the end of the LGM. Similar to Mexican *P. tremuloides* stands, the relict species *Picea mexicana* and *Picea chihuahuana* occur in isolated stands of mountainous areas in Mexico and underwent bottleneck events, respectively during the last interglacial (78-156 kyrs ago) and since 11k years ago attributed to Holocene warming (Jaramillo-Correa *et al*., 2006, 2015). Microsatellite analysis in the angiosperm *Acer saccharum* subpopulations in southern Mexico also revealed bottleneck patterns (Vargas-Rodriguez *et al*., 2015). Within the MX lineage, most SNPs were correlated to PC3; this PC displayed the highest correlation to extreme temperature followed by mean annual precipitation, which may thus be strong drivers of local adaptation within this aspen genetic lineage where large regions often experience high temperatures and severe droughts. Vicariance among isolated populations may have exerted strong selection pressures, contributing to the highest number of adaptive SNPs observed within this lineage. We initially expected the MX lineage to have a smaller effective population size than the other genetic lineages, given that MX individuals are found in isolated, high elevation stands. However, limited migration between populations within the MX lineage may have inflated the estimated effective population size at the lineage level. If we were to examine population structure on a finer scale within the MX lineage, the effective population size might in fact be much smaller (Charlesworth, 2009).

As the *between*-lineages demographic scenario 2 best explained our data, we uncovered an earliest separation between the MX and WU lineages. Our results are in line with those from Bagley *et al*. (2020), who found western coastal *P. tremuloides* populations closer related to *P. trichocarpa*, indicating a more recent shared origin in comparison to northern populations. Populations in the southern range are therefore of interest for the species’ conservation efforts, as they might encompass unique adaptive potential. The first split within the *Populus* genus (after the divergence of *Populus* and *Salix*) occurred around 46 million years ago, thereby separating the section *Abaso* (*P. mexicana*) (Wang *et al*., 2022). It is suggested that all poplars may have descended from this single ancestor in North America (Liu *et al*., 2017; Wang *et al*., 2022), and from hereon *Populus* migrated to Eurasia through the Beringia Land Bridge and further diversified (Wang *et al*., 2022; Porth *et al*., 2024b). Within section *Populus/Leuce,* several other species occur in Mexico, including *P. monticola,* Brandegee, *P. simaroa* Rzedowski and *P. guzmanantlensis* Vazquez & Cuevas (Porth *et al*., 2024b). On the other hand, the split with the Eurasian *Populus tremula*, one of the closest relatives of *P. tremuloides,* occurred around 2.3 million years ago (Wang *et al*., 2016a), corresponding to the opening of the Bering land bridge in Alaska. It would thus be of interest for future analysis to model historical population dynamics using the genetic lineages present within both species, in order to obtain more conclusive insights into the ancestral populations of *P. tremuloides.* Further phylogenetic analyses should be conducted with other poplar species functioning as outgroup, as well as test for ancient hybridisation events. The observed split between the WU and NE-NA lineages would conform to the end of the penultimate glaciation, while the last separation of the NE-NA and NW-NA lineages would correspond to the end of the LGM. We thus did not uncover evidence for an influence of a ghost refugium, such as one in Alaska. In *P. balsamifera,* a species with palaeobotanical records in Beringia, no genetic support for its presence in this region was found either, most probably due to a loss of a genetic signal resulting from drift and high migration rates from more southern populations (Breen *et al*., 2012).

The clustering of co-adapted loci, termed “islands of divergence”, have been observed in numerous species (Yeaman, 2013; Todesco *et al*., 2020; Mérot *et al*., 2021; Hämälä *et al*., 2021) and are thought to be caused by genomic rearrangements (Yeaman, 2013). For example, a study of three wild sunflower species found 37 large haplotype blocks associated with climate and soil characteristics as well as with numerous functional traits, including seed size and flowering time (Todesco *et al*., 2020). In several cases, these haplotype blocks were linked to large structural variations, in particular inversions, which provide a mechanism for supressing recombination and thereby maintaining combinations of locally adapted alleles (Todesco *et al*., 2020). A study in seaweed fly (*Coelopa frigid*) also identified non-recombining genomic regions, including putative inversions, associated with adaptive phenotypes and environmental factors (Mérot *et al*., 2021). It is unlikely that the observed clusters of SNPs in *P. tremuloides* are the result of linked neutral loci “hitchhiking” with SNPs under selection, as LD decays on average within 10 kbp in this species (Wang *et al*., 2016c). Moreover, several genes identified within the genomic region on chromosome 2 (scaffold 3) seem to be involved in abiotic stress response or development, such as homologs of the *A. thaliana* transcription factors *AtWRKY47* (Raineri *et al*., 2015; Wu *et al*., 2020) and *AtWRKY23* (Grunewald *et al*., 2012), as well as the previously identified *LACS1* gene (Goessen *et al*., 2022) that associated with individual temperature variables. *AtLACS1* acts in wax and cutin biosynthesis of the plant cuticle, an important tissue for environmental stress protection (Lü *et al*., 2009; Weng *et al*., 2010). The formation of such genomic islands of divergence could be the result of repeated cycles of divergent and adaptive selection, followed by homogenization due to environmental and demographic fluctuations, such as those observed in the alternation of ice ages and interglacial periods (Yeaman, 2013).

### Future outlook

In light of ongoing climate change affecting many North American tree species, there is an urgent need to conserve populations that are representative of the continental and regional structures of adaptive genetic diversity identified in this study. Based on our findings, we are currently conducting a study of the species’ genomic vulnerability under projected future climate conditions to identify populations of interest for conservation measures. In addition, the results from our study may help improve our understanding of the role of climate and demographic history in the survival of aspens after fires.

Overall, in our study, we examined the genomic signatures of historical population dynamics and adaptive genetic variation in a widespread species. We described how regional evolution has affected the demographic history and adaptive variation in each of the four genetic lineages of this species, which represent large geographic areas with different climate zones and vicariance factors. Genomic blocks of adaptive SNPs (islands of divergence) may act as a mechanism for maintaining local adaptation and are most likely the result of the isolation of lineages and potential secondary contacts during glacial cycles. The results of our study can serve as a blueprint for other species with similar widespread or local geographic distributions and similar life history traits.

## Supporting information

headerRF_run1

headerRF_run2

## ACKNOWLEDGEMENTS

We acknowledge funding from Fonds de recherche du Québec—Nature et technologies (FRQNT) ‘Programme bilatéral de recherche collaborative Québec-Mexique’ for Project 2018-265002 to I Porth, C Wehenkel and J Bousquet. This study was also largely supported by an NSERC Discovery Grant awarded to I Porth (RGPIN/04748-2017) as well as co-funding obtained through NRCan (N Isabel). This research was enabled by support from Digital Research Alliance of Canada to I Porth. We thank M Lamothe (NRCan) for SNP imputation script provision. We thank everyone involved in the original sample collection, including Isabel Callahan-Sánchez (student), Universidad Michoacana de San Nicolás de Hidalgo, for collecting the extreme southern *P. tremuloides* population.

## DATA ACCESSIBILITY AND BENEFIT-SHARING

Raw sequence data and related metadata are deposited in the Sequence Read Archive (BioProject ID PRJNA809939) and will be made available upon publication. Benefits Generated: The benefits of this research result from the sharing of our data and findings with the broader scientific community. The provisions of the Nagoya protocol on Access and Benefit-sharing were complied with during sampling and transfer of plant material from Mexico into Canada: The Mexican project partners had access to all Mexican genetic data for *P. tremuloides*, as specified in the funded project; the permits for the transfer of plant material were obtained in accordance with the Canadian Food Inspection Agency for import into Canada, and the Mexican project partners received export permits; The results of this project will be used to conserve *P. tremuloides* populations in the Mexican mountain ranges by identifying vulnerable stands and thus preserving biodiversity.

## AUTHOR CONTRIBUTIONS

IP, RG designed the overall study. NI, IP, CW, AB, JV, EM, CS, RG provided samples in the experimental study design. RG wrote the manuscript and performed all analyses. IP, JB, NI helped with complementary funding of the study and revisions of manuscript drafts. All authors read and approved the final manuscript.

## SUPPORTING INFORMATION

**Figure S1.** Representation of the tested demographic scenarios using fastsimcoal2.

**Figure S2.** All 10 tested scenarios between genetic lineages using DIY-ABC-RF for Run1.

**Figure S3.** All 13 tested scenarios between genetic lineages using DIY-ABC-RF for Run2.

**Figure S4.** Projection of the training set data on the first two linear discriminant analysis (LDA) axes when analysing all 10 scenarios of Run1.

**Figure S5.** Projection of the training set data on the first two linear discriminant analysis (LDA) axes when analysing all 13 scenarios of Run2.

**Figure S6 a-b.** Rangewide Weir and Cockerham *F*_ST_ heatmap and maximum likelihood tree.

**Figure S7 a-f.** Principal component (PC) 1 and PC2 using 24 environmental variables available from the climateNA database (Wang *et al*., 2016a).

**Figure S8 a-f.** Principal component (PC) 1 and PC3 using 24 environmental variables available from the climateNA database (Wang *et al*., 2016a).

**Figure S9.** Correlation analysis of all environmental variables.

**Figure S10.** Correlation analysis of selected non-collinear environmental variables.

**Figure S11 a-b.** PC1 vs. PC2 for all environmental variables and individuals for the dataset filtered with MAS=15.

**Figure S12 a-b.** PC1 vs. PC3 for all environmental variables and individuals for the dataset filtered with MAS=15.

**Figure S13 a-f.** Manhattan plots of results for Bayenv2 (Bayes factor, figs. a-c) and LFMM2 (adjusted *P* value (*P*), figs. d-f) for PC1 (figs. a, d), PC2 (figs. b, e) and PC3 (figs. c, f) for all samples along the 20 major scaffolds using a minor allele count (MAS) filter of 15.

**Figure S14 a-c.** Manhattan plots of results for Bayenv2 for PC1 (fig. a), PC2 (fig. b) and PC3 (fig. c) calculated over selected non-collinear environmental variables (MWMT, MCMT, MAP, MSP, FFP, AHM, PAS, EXT, RH and DD1040) for all samples along the 20 major scaffolds with a minor allele count (MAS) filter of 15.

**Figure S15.** Overlap of identified SNPs between the original bayenv2 and LFMM2 analyses (with MAS = 56 filter), vs. the new bayenv2 and LFMM2 analyses (with MAS = 15 filter) on PC1 calculated over all selected variables.

**Figure S16.** Overlap of identified SNPs between the original bayenv2 and LFMM2 analyses (with MAS = 56 filter), vs. the new bayenv2 and LFMM2 analyses (with MAS = 15 filter) on PC2 calculated over all selected variables.

**Figure S17.** Overlap of identified SNPs between the original bayenv2 and LFMM2 analyses (with MAS = 56 filter), vs. the new bayenv2 and LFMM2 analyses (with MAS = 15 filter) on PC3 calculated over all selected variables.

**Figure S18.** Linkage-disequilibrium (LD, r^2^) plots of SNPs identified in GEA analyses for climate PC1, PC2 and PC3 using Bayenv2 and LFMM2 (rangewide scale).

**Figure S19.** Classification of genes containing the SNPs or being near the SNPs that were associated with environmental PC1, PC2 and PC3 identified within each genetic lineage.

**Figure S20.** Frequency of PFAMs and SUPERFAMILYs for genes with or near SNPs identified for PC1, PC2 and PC3 within each genetic lineage or over the entire sample size.

**Table S1.** Comparison of SNP calling with old and new reference genome without any filtering.

**Table S2.** Filtering settings for dataset Set_01 (used for *F*_ST_, phylogeny and historical gene flow analyses) and Set_02 (used for genotype-environment association (GEA) analyses).

**Table S3.** Abbreviations and explanations of environmental variables of ClimateNA database.

**Table S4.** Statistical results for best model runs of demographic scenarios tested within single lineages with fastsimcoal2 using “bounded” keyword for upper range in time parameter (max. 2,000 years).

**Table S5.** Scenario prediction votes of Run1 using a forest of 1500 trees, in DIY-ABC-RF.

**Table S6.** Scenario prediction votes of Run2 using a forest of 1500 trees, in DIY-ABC-RF.

**Table S7.** The contingency table of true vs. predicted scenarios for each sample in the training set for Run01, generated by DIY-ABC-RF.

**Table S8.** The contingency table of true vs. predicted scenarios for each sample in the training set for Run02, generated by DIY-ABC-RF.

**Table S9.** Number of identified environmentally associated SNPs per scaffold for all 24 environmental variables and PC1, PC2 and PC3, based on the entire sample size.

**Table S10.** Number of overlapping SNPs between the original runs (Bayenv and LFMM with MAS=56 filter) vs. the new runs (Bayenv and LFMM with MAS=15 filter) for PC1-2-3 calculated on all environmental variables.

**Table S11.** Number of SNPs and genes identified with LFMM2 and Bayenv2 per tested climatic variable within each genetic lineage.

**Methods S1.** Comparison of SNP calling using either the old or a new *Populus tremuloides* reference genome.

**Methods S2.** Fastsimcoal2 to infer demographic scenarios for each genetic lineage.

**Methods S3.** DIY-ABC-RF to deduce the demographic history between genetic lineages.

**Methods S4.** Genotype-environment association analysis on a dataset with relaxed filtering (for MAS) and selection of environmental variables based on correlation.

**Methods S5.** Genotype-environment association analyses methods.

**Results 1.** Genotype-environment association analyses with the relaxed filtering.

**Dataset S1.** Overview of sample collection in this study. This includes individual names, population names, geographical coordinates, species, library number, and ploidy assignment.

**Dataset S2.** Overview of climate variables used in the genotype-environment associations.

**Dataset S3.** SNPs identified for all environmental variables using Bayenv2 over the entire sample size. Explanation of column names are provided in the spreadsheet.

**Dataset S4.** SNPs identified for all environmental variables using LFMM2 over the entire sample size. Explanation of column names are given in the spreadsheet.

**Dataset S5.** SNPs identified for PC1, PC2 and PC3 using Bayenv2 and LFMM2 over the entire sample size. Explanation of column names are provided within the spreadsheet.

**Dataset S6.** Inversion data of four genetically distinct individuals.

**Dataset S7.** SNPs identified for PC1, PC2 and PC3 using Bayenv2 and LFMM2 within each respective genetic lineage. Explanation of column names are found in the spreadsheet.

**Dataset S8.** Annotation summary of SNPs identified for PC1, PC2 and PC3 using Bayenv2 and LFMM2 within each respective lineage.

**Dataset S9.** SNPs identified for all environmental variables using LFMM2 within each respective lineage. Explanation of column names are given within the spreadsheet.

**headerRF_run1.txt** The input file for DIY-ABC-RF2 for Run1. The file described all tested scenarios and input parameters.

**headerRF_run2.txt** The input file for DIY-ABC-RF2 for Run2. The file described all tested scenarios and input parameters.

