## Supplemental file for "Elucidating continental-wide phylogeographic and adaptive processes shaping the genome-wide diversity of North America’s most widely distributed tree"

The following Supporting Information is available for this article:

**Figure S1.** Representation of the tested demographic scenarios using fastsimcoal2.

**Figure S2.** All 10 tested scenarios between genetic lineages using DIY-ABC-RF for Run1.

**Table S10.** Number of overlapping SNPs between the original runs (Bayenv2 and LFMM2 with MAS=56 filter) vs. the new runs (Bayenv2 and LFMM2 with MAS=15 filter) for PC1-2-3 calculated on all environmental variables.

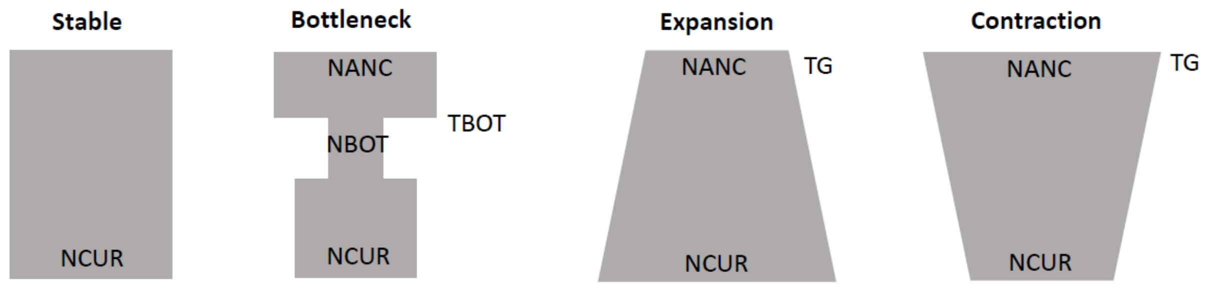


**Figure S1. Representation of the tested demographic scenarios using fastsimcoal2.** Stable, bottleneck, expansion and contraction scenarios were tested within each genetic lineage, *i.e.* northeast North America (NE-NA), northwest North America (NW-NA), western US (WU) and Mexico (MX). NANC indicates ancient effective population size, NCUR indicates current effective population size, TBOT indicates start of bottleneck in generations, NBOT indicates effective population size during bottleneck, TG indicates the number of generations since expansion or contraction.

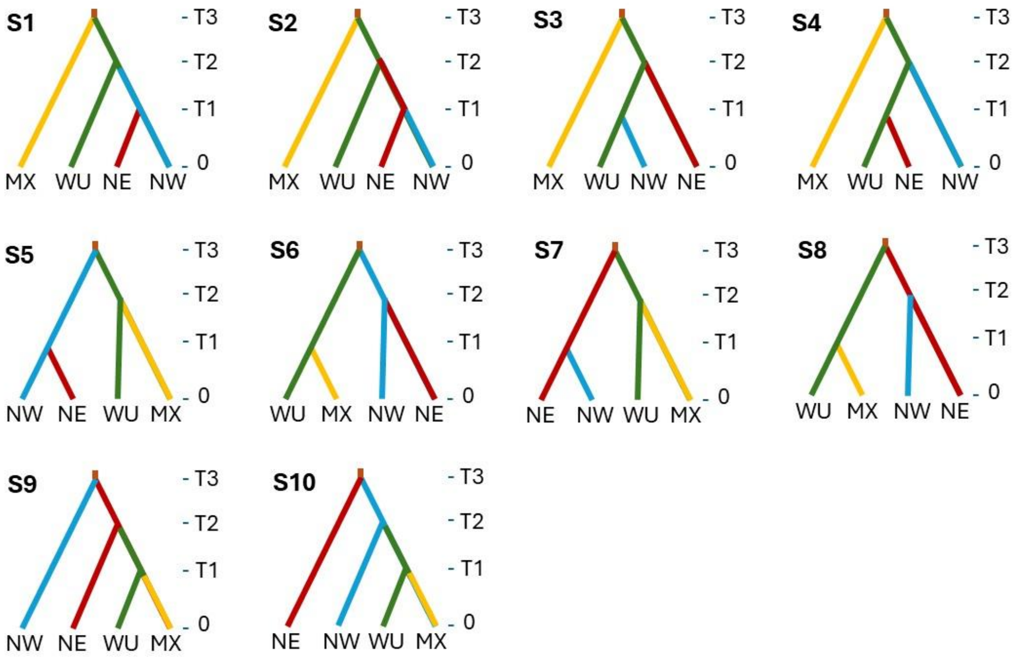


**Figure S2. All 10 tested scenarios between genetic lineages using DIY-ABC-RF for Run1.**

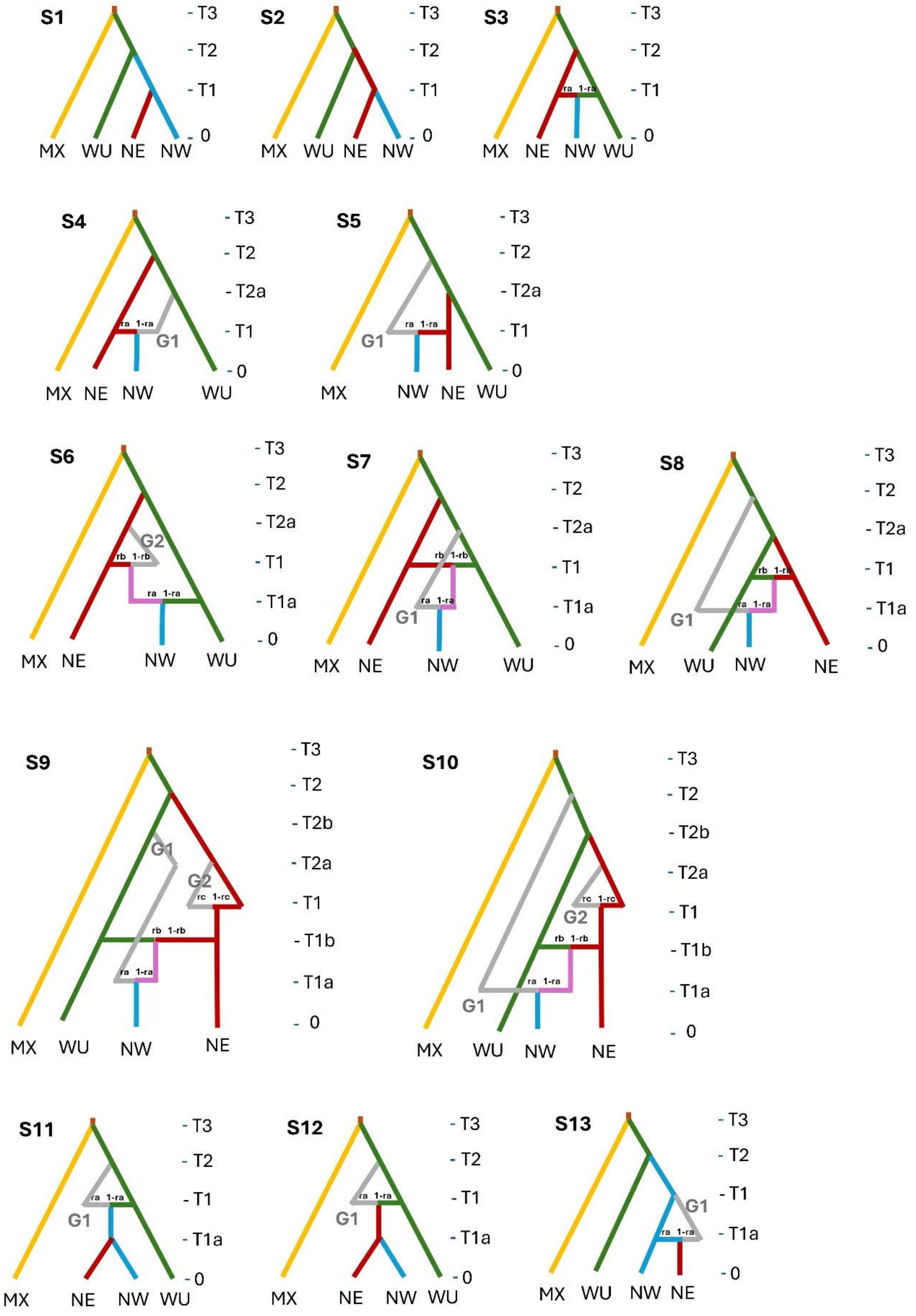


**Figure S3. All 13 tested scenarios between genetic lineages using DIY-ABC-RF for Run2.**

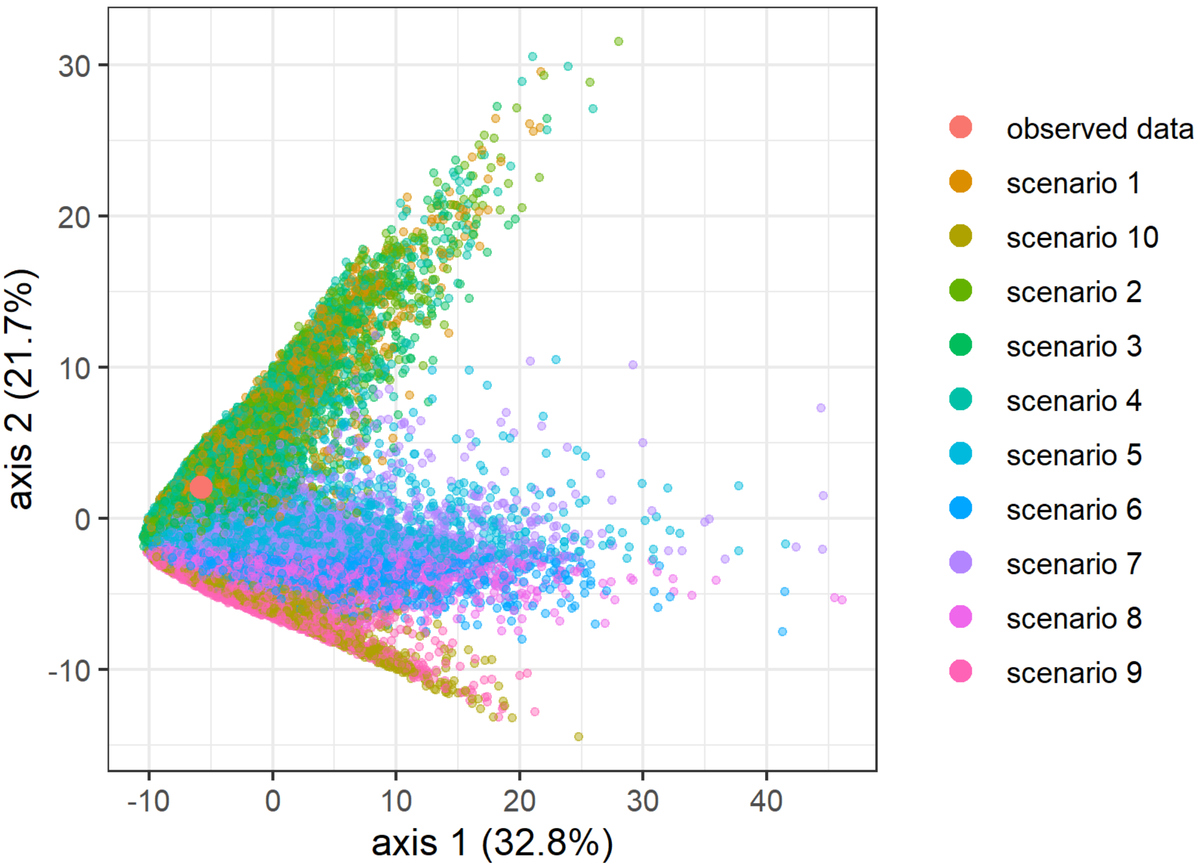


**Figure S4. Projection of the training set data on the first two linear discriminant analysis (LDA) axes when analysing all 10 scenarios of Run1.** The observed dataset is indicated with the large pink dot. The figure is generated by DIY-ABC-RF software.

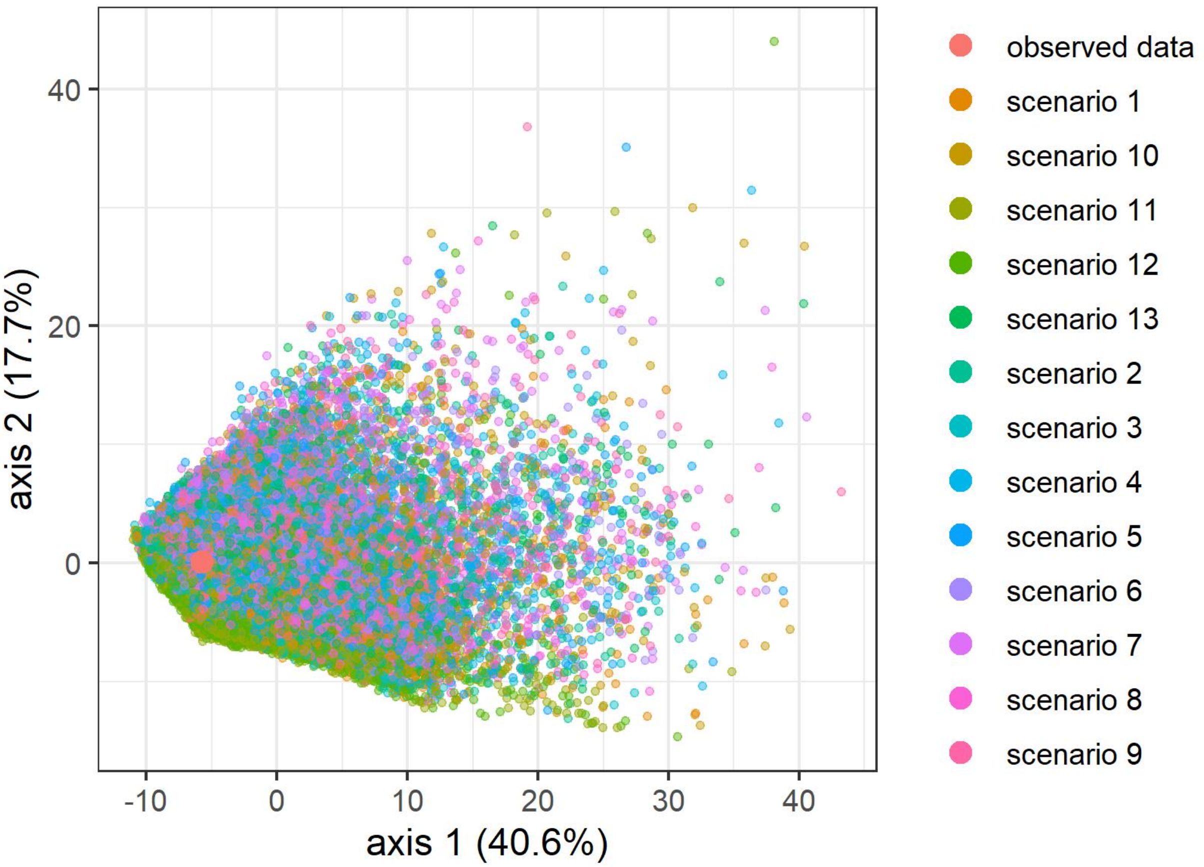


**Figure S5. Projection of the training set data on the first two linear discriminant analysis (LDA) axes when analysing all 13 scenarios of Run2.** The observed dataset is indicated with the large pink dot. The figure is generated by DIY-ABC-RF software.

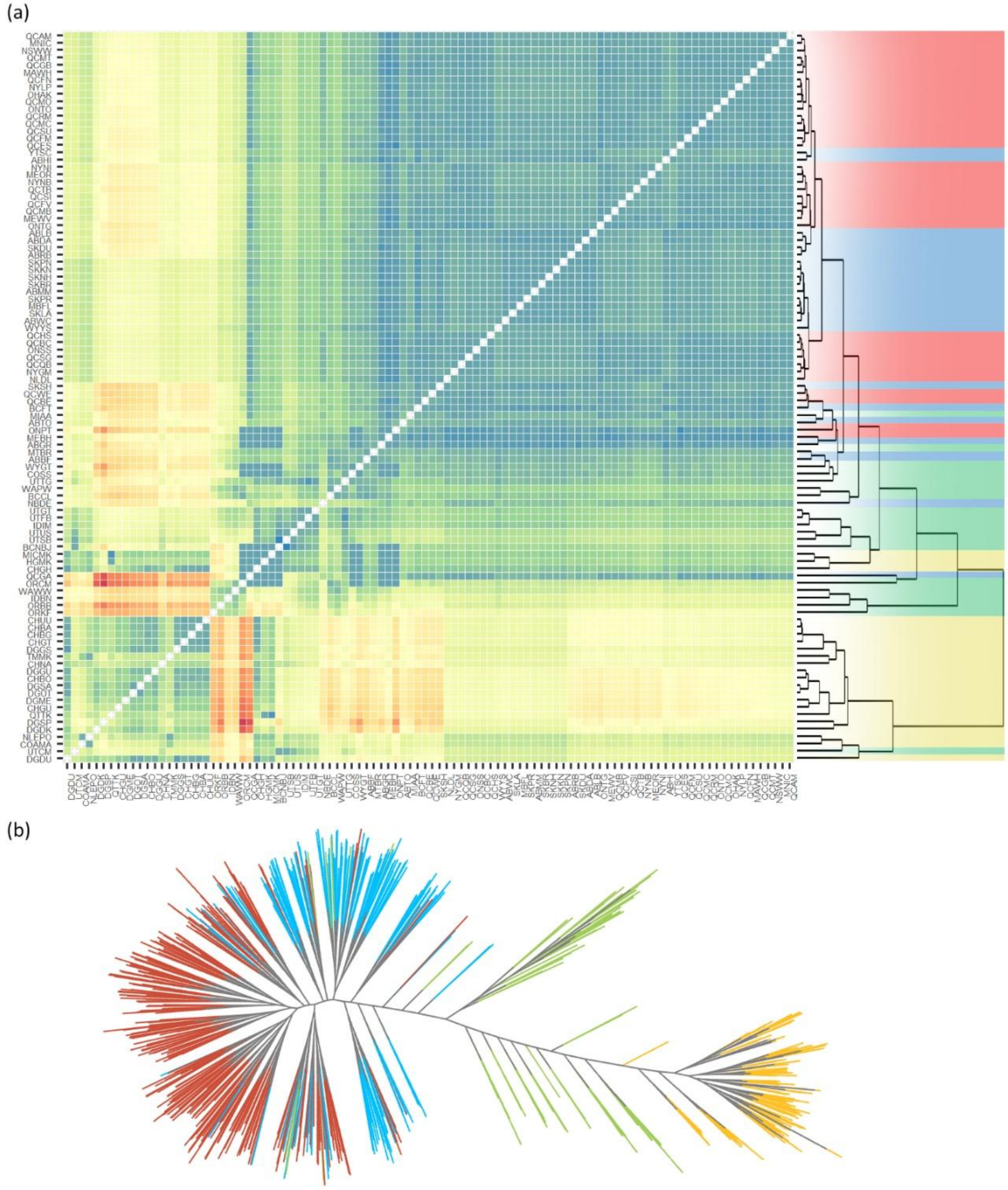


**Figure S6 a-b. Rangewide Weir and Cockerham $F_{ST}$ heatmap and maximum likelihood tree.** **a.** $F_{ST}$ heatmap with colors shading underlying the UPGMA tree (right) indicate lineage as determined by Goessen *et al*. (2022). Red indicates NE-NA, blue NW-NA, green WU, yellow MX genetic lineages. The color chart on the right indicates the $F_{ST}$ values of the heatmap, with red indicating higher values and blue indicating lower values. The analyses were performed on a population basis. **b.** Maximum likelihood tree generated by RaxML and visualized with ggtree in R with "equal angle" layout. Tree was generated with 6000 random SNPs from Set_01. Dot colors represent the origin of the four genetic lineages: red for NE-NA blue for NW-NA, green for WU and yellow for MX. The analysis was performed on an individual basis.

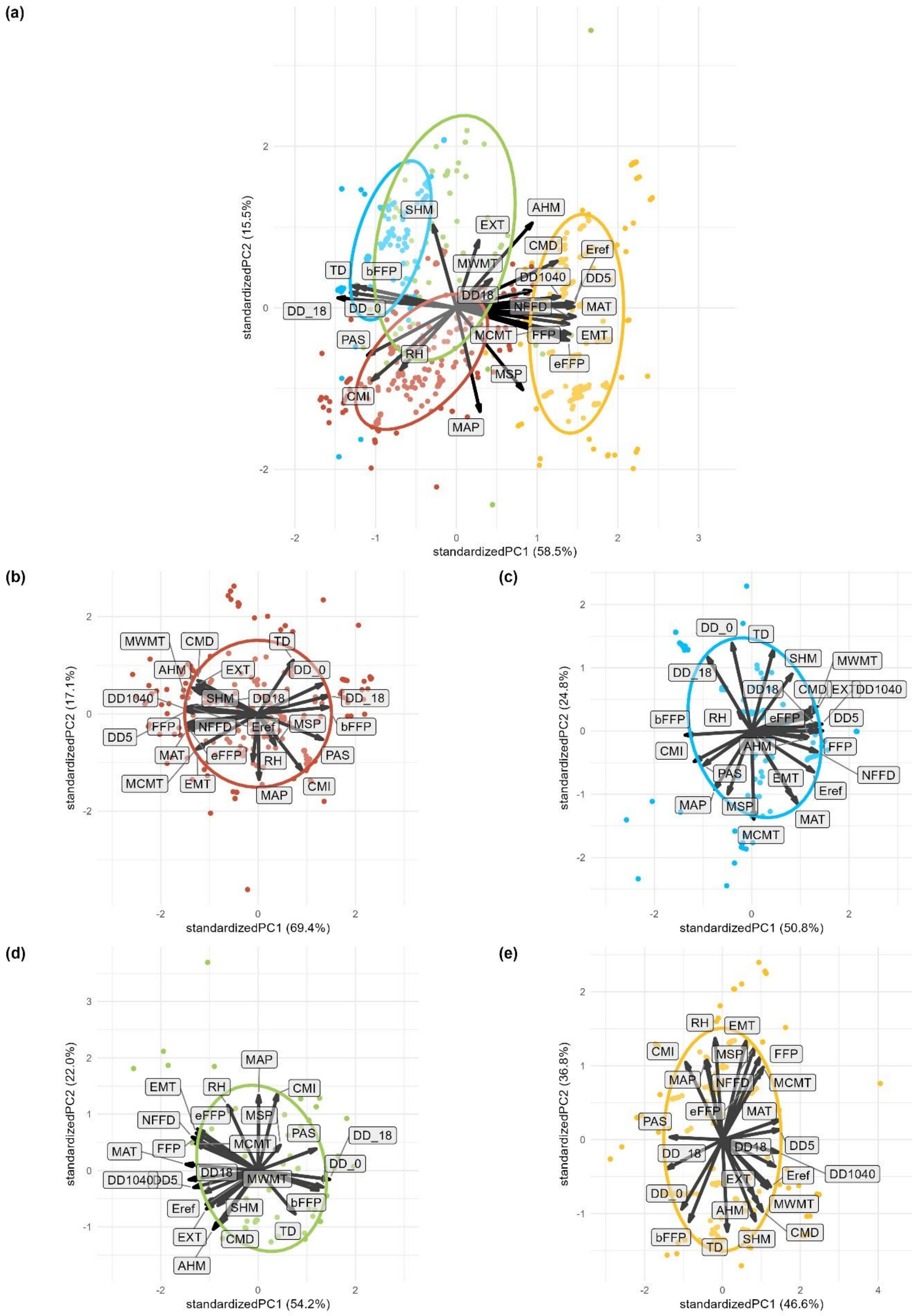
(a)
standardizedPC2 (15.5%)
standardizedPC1 (58.5%)
SHM
AHM
EXT
CMD
Eref
TD
bFFP
MWMT
DD1040
DD5
DD18
DD_18
DD_0
NFFD
MAT
MCMT
FFP
EMT
PAS
RH
eFFP
MSP
CMI
MAP
(b)
standardizedPC2 (17.1%)
standardizedPC1 (69.4%)
MWMT
CMD
TD
AHM
EXT
DD_0
DD1040
SHM
DD18
DD_18
FFP
NFFD
Eref
MSP
bFFP
DD5
MAT
eFFP
RH
PAS
MCMT
EMT
CMI
MAP
(c)
standardizedPC2 (24.8%)
standardizedPC1 (50.8%)
DD_0
TD
SHM
MWMT
DD_18
DD18
CMD
EXT
DD1040
bFFP
RH
eFFP
DD5
AHM
FFP
CMI
NFFD
PAS
EMT
Eref
MAP
MSP
MAT
MCMT
(d)
standardizedPC2 (22.0%)
standardizedPC1 (54.2%)
MAP
EMT
RH
CMI
NFFD
eFFP
MSP
PAS
DD_18
MAT
FFP
MCMT
DD_0
DD18
DD1040
DD5
MWMT
bFFP
Eref
SHM
EXT
CMD
TD
AHM
(e)
standardizedPC2 (36.8%)
standardizedPC1 (46.6%)
RH
EMT
CMI
MSP
FFP
MAP
NFFD
MCMT
eFFP
MAT
PAS
DD_18
DD18
DD5
Eref
DD1040
EXT
DD_0
AHM
MWMT
CMD
bFFP
TD
SHM

**Figure S7 a-e. Principal component (PC) 1 and PC2 using 24 environmental variables available from the climateNA database (Wang *et al.*, 2016b). a.** All samples, with colors indicating the origin of lineages, with red for northeast North America (NE-NA), blue for northwest North America (NW-NA), green for western US (WU) and yellow for the Mexico (MX). **b.** Samples within the NE-NA lineage. **c.** Samples within the NW-NA lineage. **d.** Samples within the WU lineage. **e.** Samples within the MX lineage. Ellipses represent the default 68% normal probability. Climatic variables include: mean annual temperature (MAT, °C), mean warmest month temperature (MWMT, °C), mean coldest month temperature (MCMT, °C), temperature difference between MWMT and MCMT (TD, °C), mean annual precipitation (MAP, mm), May to September precipitation (MSP, mm), annual heat-moisture index (AHM, (MAT+10)/(MAP/1000)), summer heat-moisture index (SHM, (MWMT/(MSP/1000)), degree-days below 0°C (DD_0), degree-days above 5°C (DD5), degree-days below 18°C (DD_18), degree-days above 18°C (DD18), number of frost-free days (NFFD), frost-free period (FFP), the day of the year on which FFP begins (bFFP), the day of the year on which FFP ends (eFFP), precipitation as snow (PAS, mm), extreme minimum temperature over 30 years (EMT), extreme maximum temperature over 30 years (EXT), Hargreaves reference evaporation (Eref, mm), Hargreaves climatic moisture deficit (CMD, mm), mean annual relative humidity (RH, %), Hogg's climate moisture index (CMI, mm), degree-days above 10°C and below 40°C (DD1040).

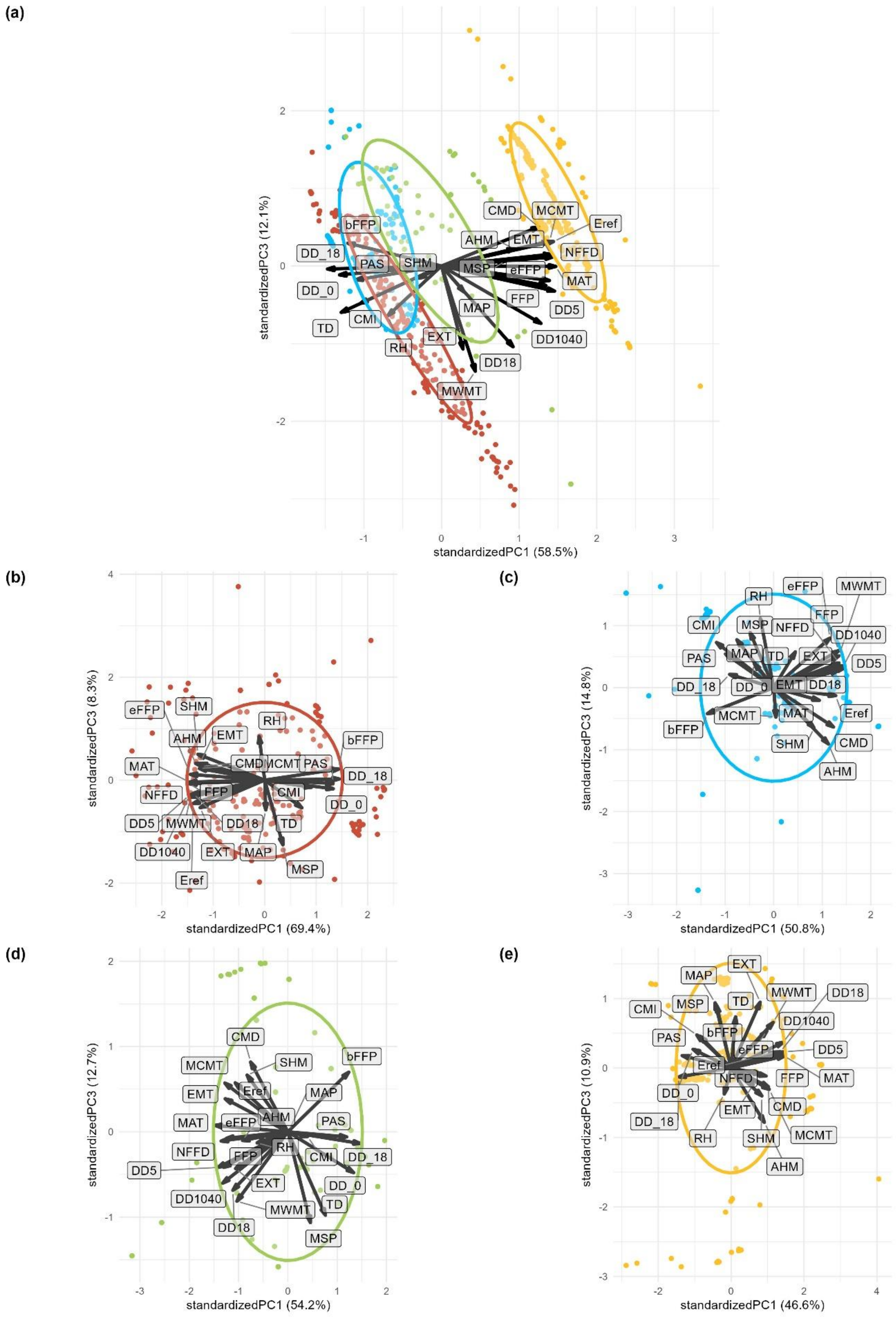
(a)
standardizedPC1 (58.5%)
standardizedPC3 (12.1%)
CMD
MCMT
Eref
bFFP
AHM
EMT
NFFD
DD_18
PAS
SHM
MSP
eFFP
MAT
DD_0
FFP
DD5
MAP
CMI
TD
EXT
DD1040
RH
DD18
MWMT
(b)
standardizedPC1 (69.4%)
standardizedPC3 (8.3%)
eFFP
SHM
RH
bFFP
AHM
EMT
MAT
CMD
MCMT
PAS
DD_18
FFP
CMI
NFFD
DD_0
DD5
MWMT
DD18
TD
DD1040
EXT
MAP
MSP
Eref
(c)
standardizedPC1 (50.8%)
standardizedPC3 (14.8%)
RH
eFFP
MWMT
FFP
CMI
MSP
NFFD
DD1040
PAS
MAP
TD
EXT
DD5
DD_18
DD_0
EMT
DD18
MCMT
MAT
Eref
bFFP
SHM
CMD
AHM
(d)
standardizedPC1 (54.2%)
standardizedPC3 (12.7%)
CMD
SHM
bFFP
MCMT
Eref
MAP
EMT
MAT
eFFP
AHM
PAS
NFFD
FFP
RH
CMI
DD_18
DD5
EXT
DD_0
DD1040
MWMT
TD
DD18
MSP
(e)
standardizedPC1 (46.6%)
standardizedPC3 (10.9%)
EXT
MAP
MWMT
DD18
MSP
TD
CMI
DD1040
bFFP
PAS
eFFP
DD5
Eref
NFFD
FFP
MAT
DD_0
CMD
EMT
DD_18
MCMT
RH
SHM
AHM

**Figure S8 a-e. Principal component (PC) 1 and PC3 using 24 environmental variables available from the climateNA database (Wang *et al.*, 2016b). a.** All samples, with colors indicating the origins of genetic lineages, with red for northeast North America (NE-NA), blue for northwest North America (NW-NA), green for western US (WU) and yellow for Mexico (MX). **b.** Samples within the NE-NA lineage. **c.** Samples within the NW-NA lineage. **d.** Samples within the WU lineage. **e.** Samples within the MX lineage. Ellipses represent the default 68% normal probability. Climatic variables include: mean annual temperature (MAT, °C), mean warmest month temperature (MWMT, °C), mean coldest month temperature (MCMT, °C), temperature difference between MWMT and MCMT (TD, °C), mean annual precipitation (MAP, mm), May to September precipitation (MSP, mm), annual heat-moisture index (AHM, (MAT+10)/(MAP/1000)), summer heat-moisture index (SHM, (MWMT/(MSP/1000)), degree-days below 0°C (DD_0), degree-days above 5°C (DD5), degree-days below 18°C (DD_18), degree-days above 18°C (DD18), number of frost-free days (NFFD), frost-free period (FFP), the day of the year on which FFP begins (bFFP), the day of the year on which FFP ends (eFFP), precipitation as snow (PAS, mm), extreme minimum temperature over 30 years (EMT), extreme maximum temperature over 30 years (EXT), Hargreaves reference evaporation (Eref, mm), Hargreaves climatic moisture deficit (CMD, mm), mean annual relative humidity (RH, %), Hogg's climate moisture index (CMI, mm), degree-days above 10°C and below 40°C (DD1040).

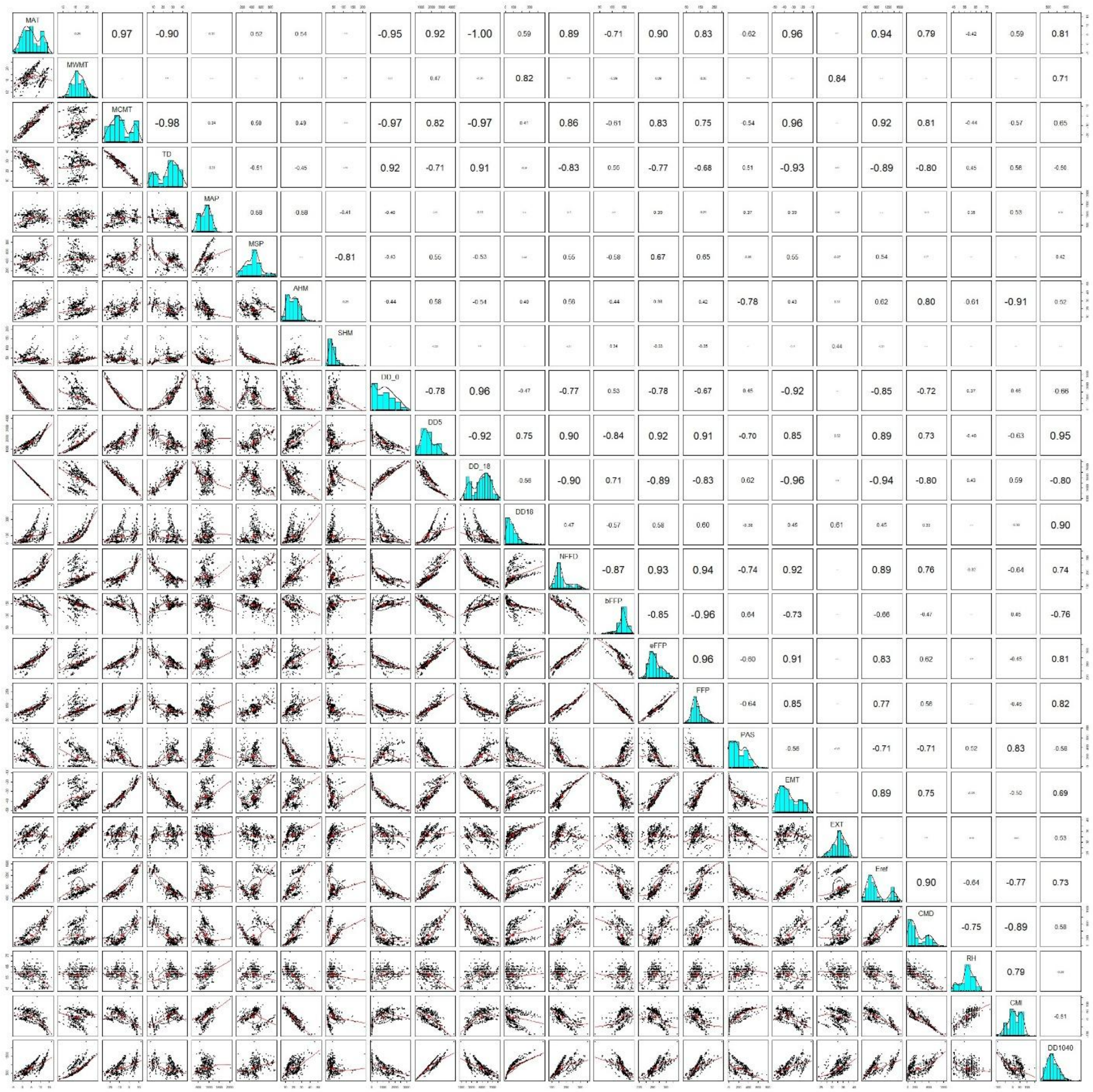

**Figure S9. Correlation analysis of all environmental variables.** Number size indicates correlation strength, with larger numbers having higher absolute correlation.

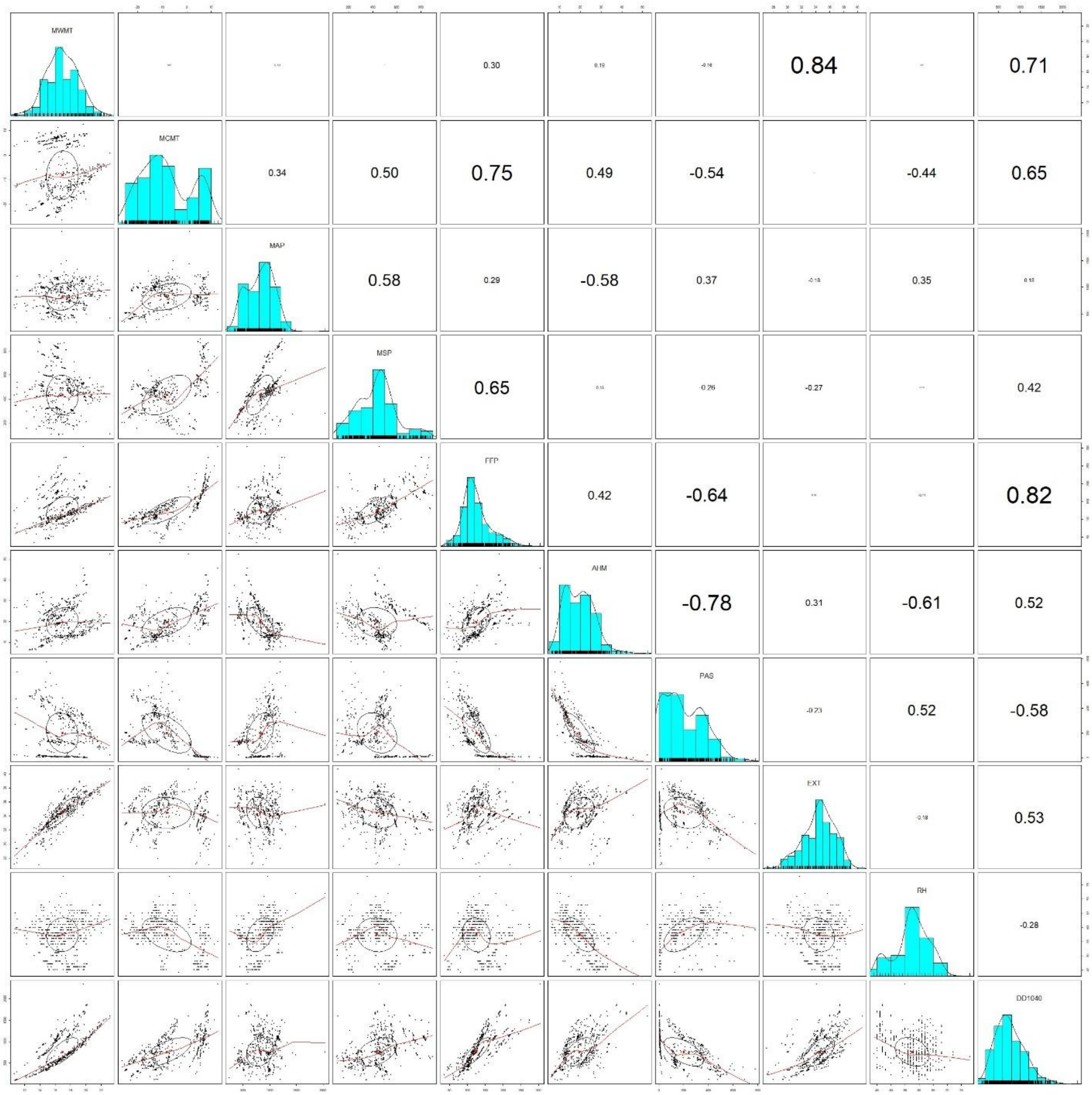


**Figure S10. Correlation analysis of selected non-collinear environmental variables.**

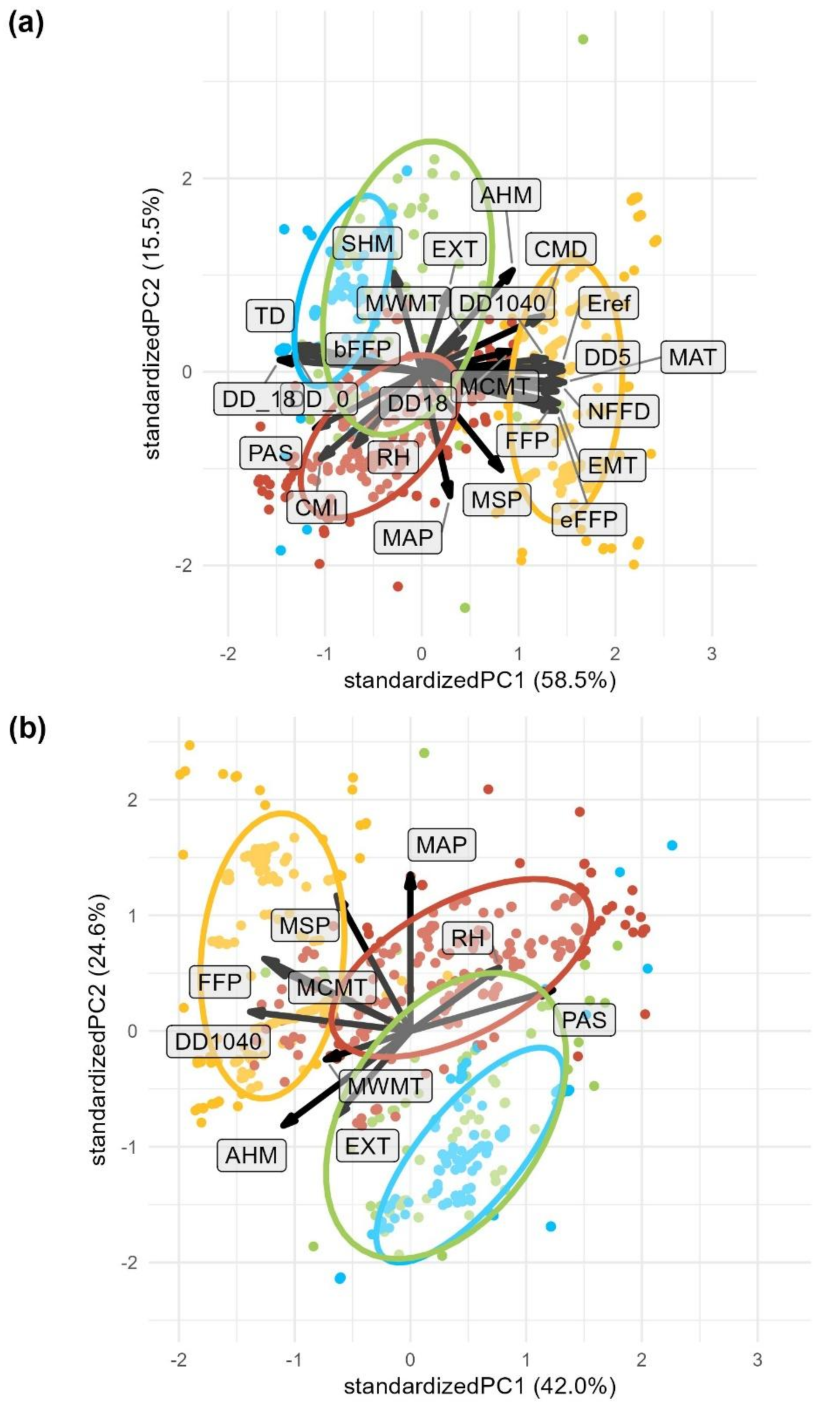


**Figure S11 a-b. PC1 vs. PC2 for all environmental variables and individuals for the dataset filtered with MAS=15**. **a.** For all variables. **b.** For selected non-collinear variables MWMT, MCMT, MAP, MSP, FFP, AHM, PAS, EXT, RH and DD1040.

(a)

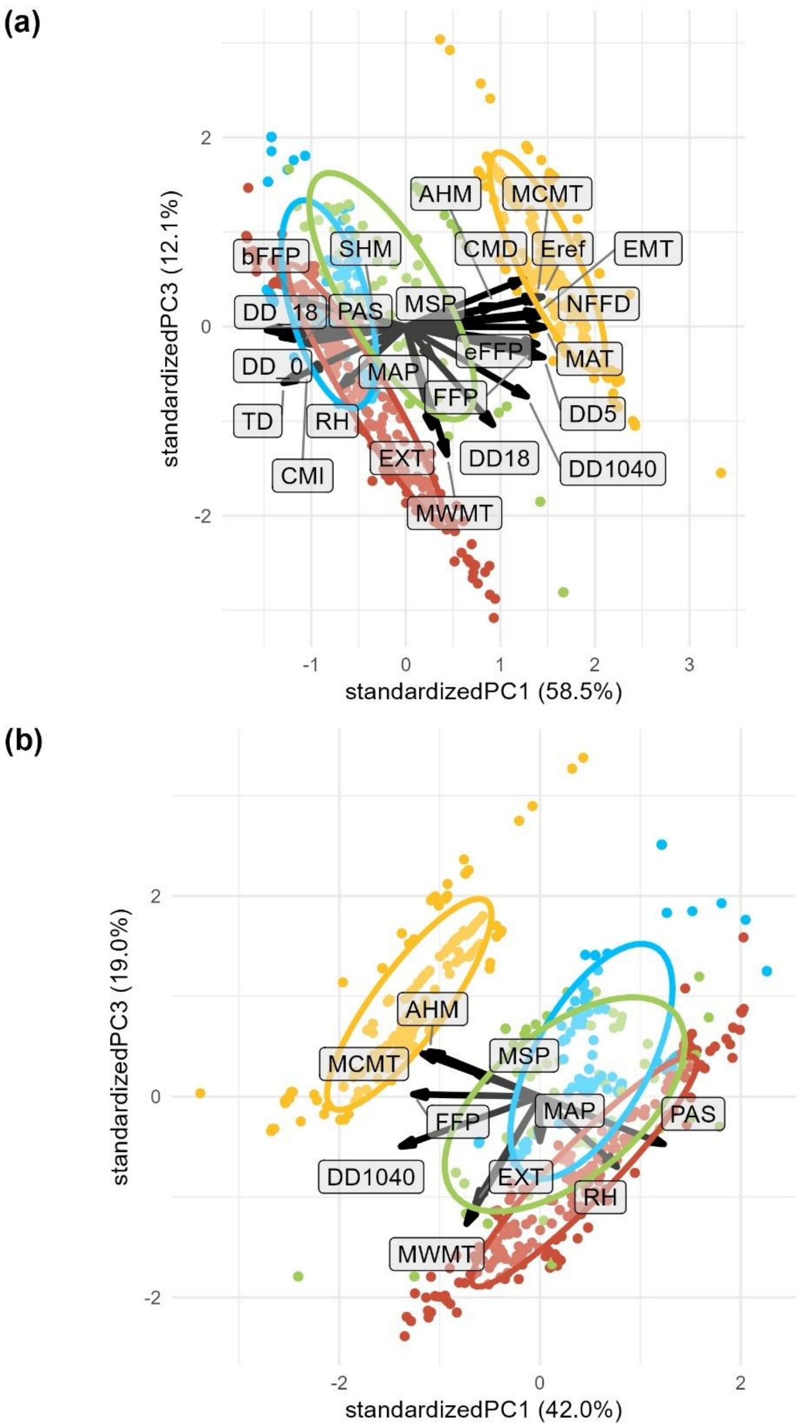


(b)

**Figure S12 a-b. PC1 vs. PC3 for all environmental variables and individuals for the dataset filtered with MAS=15**. **a.** For all variables. **b.** For selected non-collinear variables MWMT, MCMT, MAP, MSP, FFP, AHM, PAS, EXT, RH and DD1040.

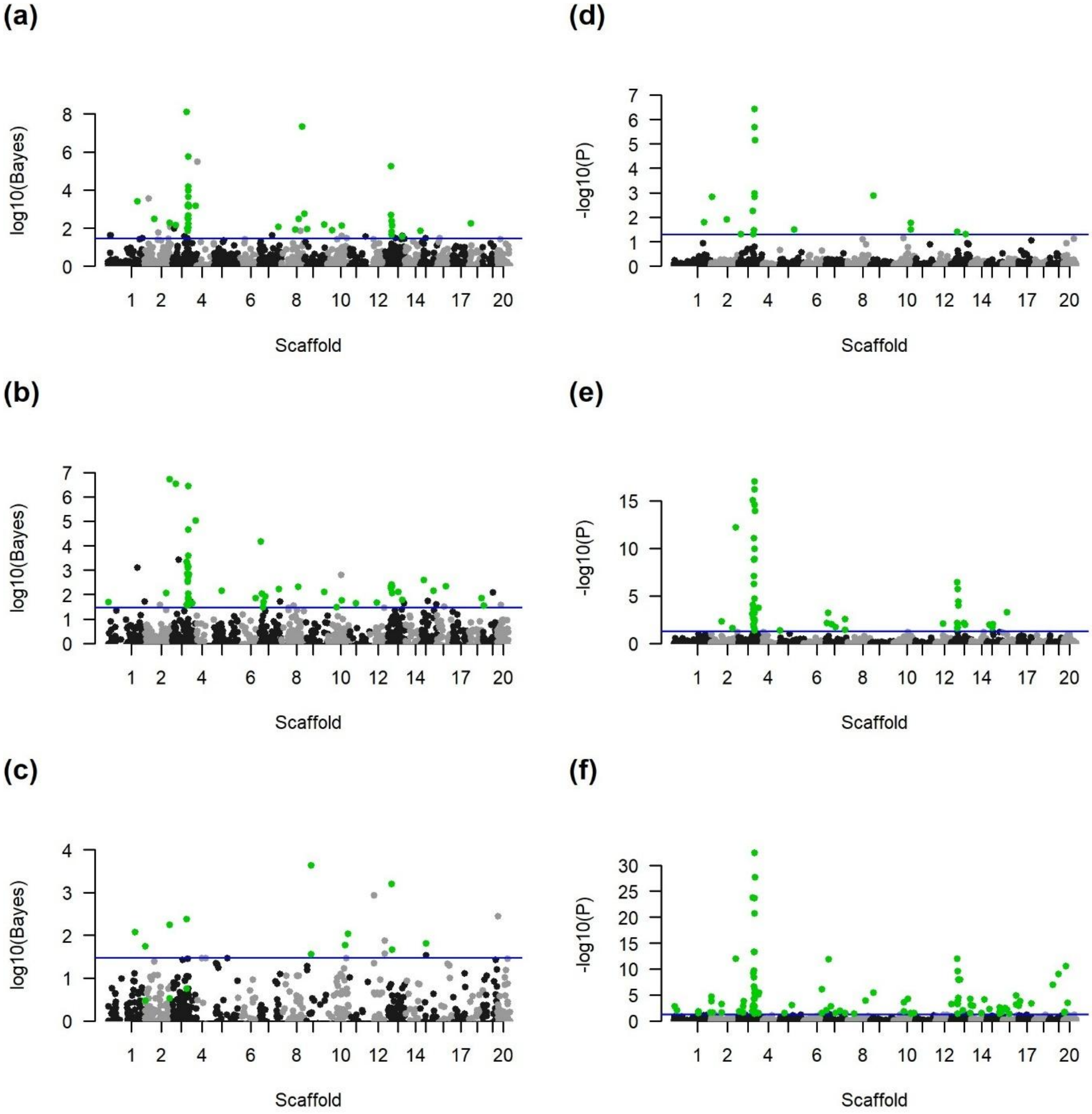


**Figure S13a-f. Manhattan plots of results for Bayenv2 (Bayes factor, figs. a-c) and LFMM2 (adjusted *P* value (*P*), figs. d-f) for PC1 (figs. a, d), PC2 (figs. b, e) and PC3 (figs. c, f) for all samples along the 20 major scaffolds using a minor allele count (MAS) filter of 15.** The blue line for Bayenv2 plots indicates the cut-off Bayes factor of 30. The blue line for LFMM2 indicates the adjusted *P* value cut-off of 0.05. SNPs identified by Bayenv2 that are above the blue cut-off line, but not colored green (Figs. a-c) did not pass the criteria for overlap with the top 5% for Pearson and Spearman correlations.

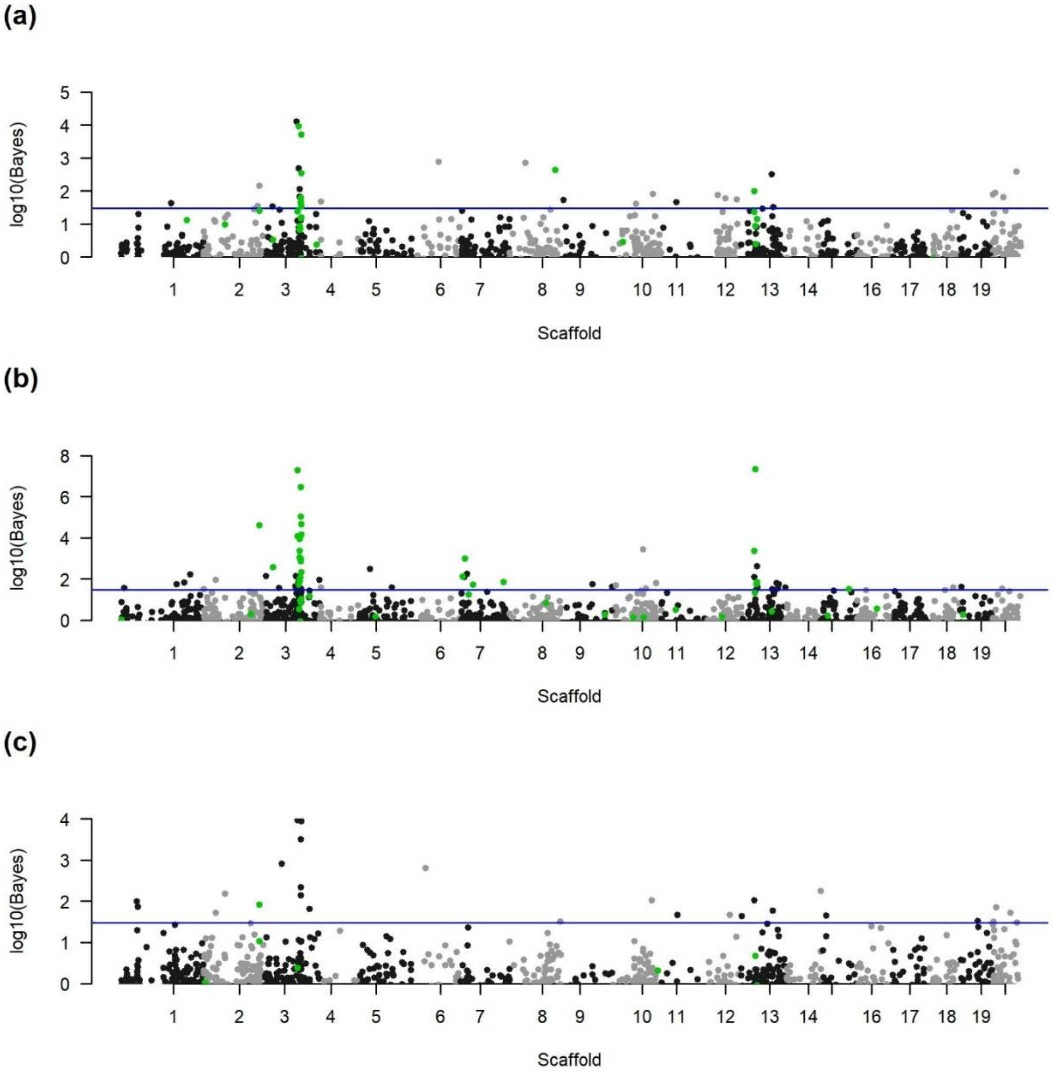


**Figure S14 a-c. Manhattan plots of results for Bayenv2 for PC1 (fig. a), PC2 (fig. b) and PC3 (fig. c) calculated over selected non-collinear environmental variables (MWMT, MCMT, MAP, MSP, FFP, AHM, PAS, EXT, RH and DD1040) for all samples along the 20 major scaffolds with a minor allele count (MAS) filter of 15.** The blue line for Bayenv2 plots indicates the cut-off Bayes factor of 30. SNPs identified by Bayenv2 that are above the blue cut-off line, but not colored green did not pass the criteria for overlap with the top 5% for Pearson and Spearman correlations.

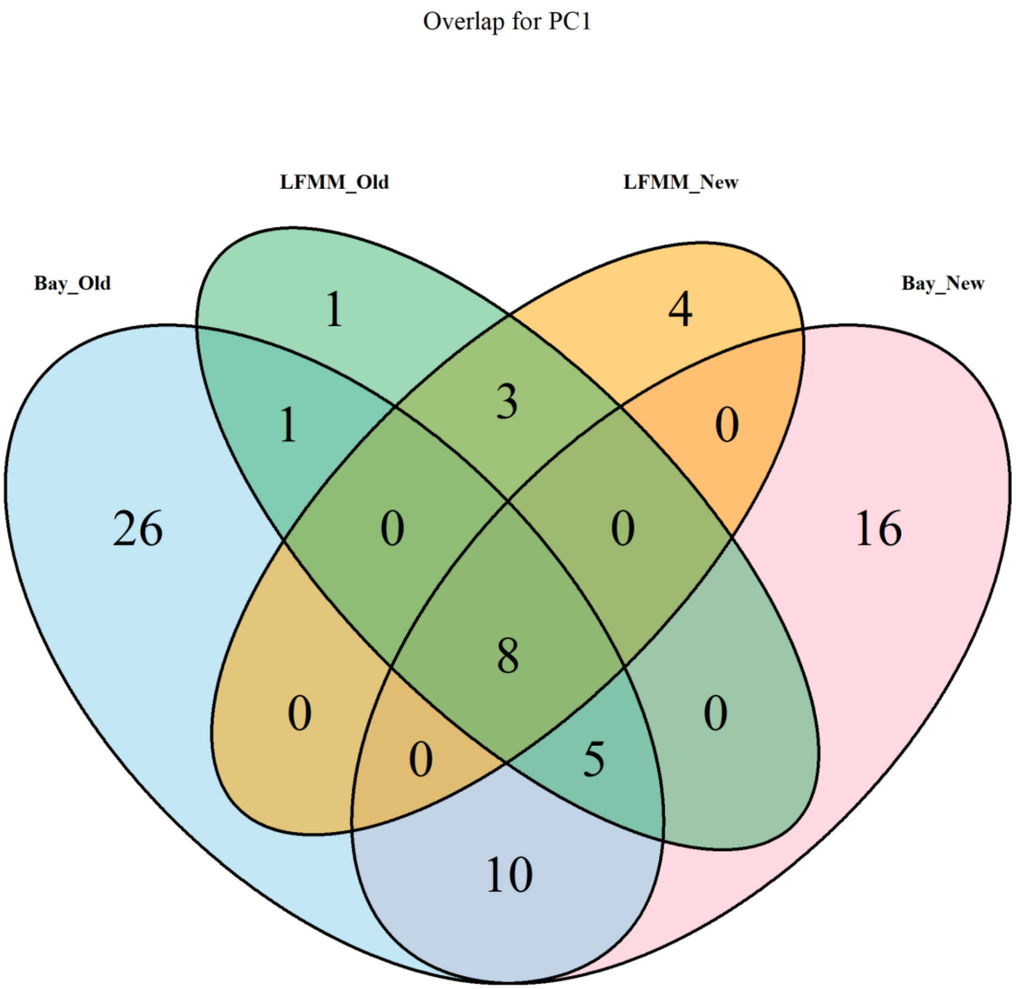


**Figure S15. Overlap of identified SNPs between the original bayenv2 and LFMM2 analyses (with MAS = 56 filter), vs. the new bayenv2 and LFMM2 analyses (with MAS = 15 filter) on PC1 calculated over all selected variables.**

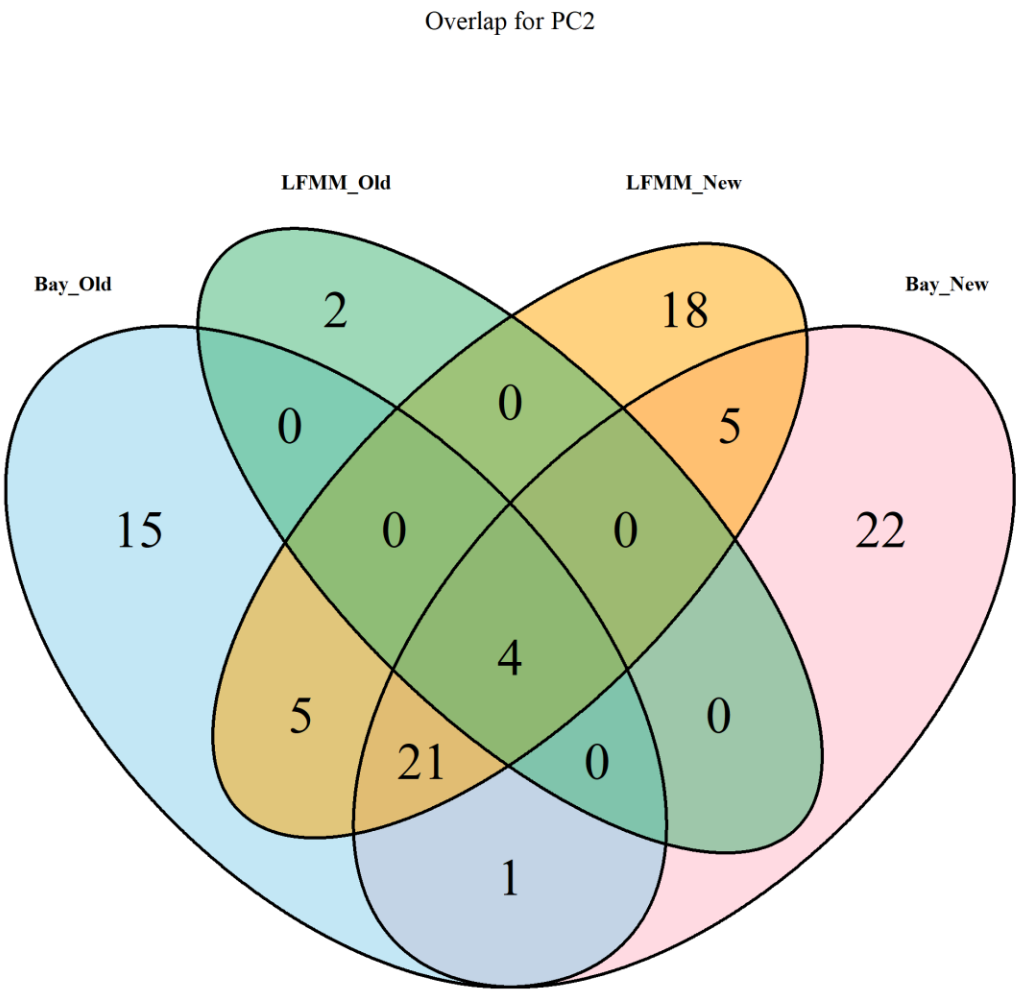


**Figure S16. Overlap of identified SNPs between the original bayenv2 and LFMM2 analyses (with MAS = 56 filter), vs. the new bayenv2 and LFMM2 analyses (with MAS = 15 filter) on PC2 calculated over all selected variables.**

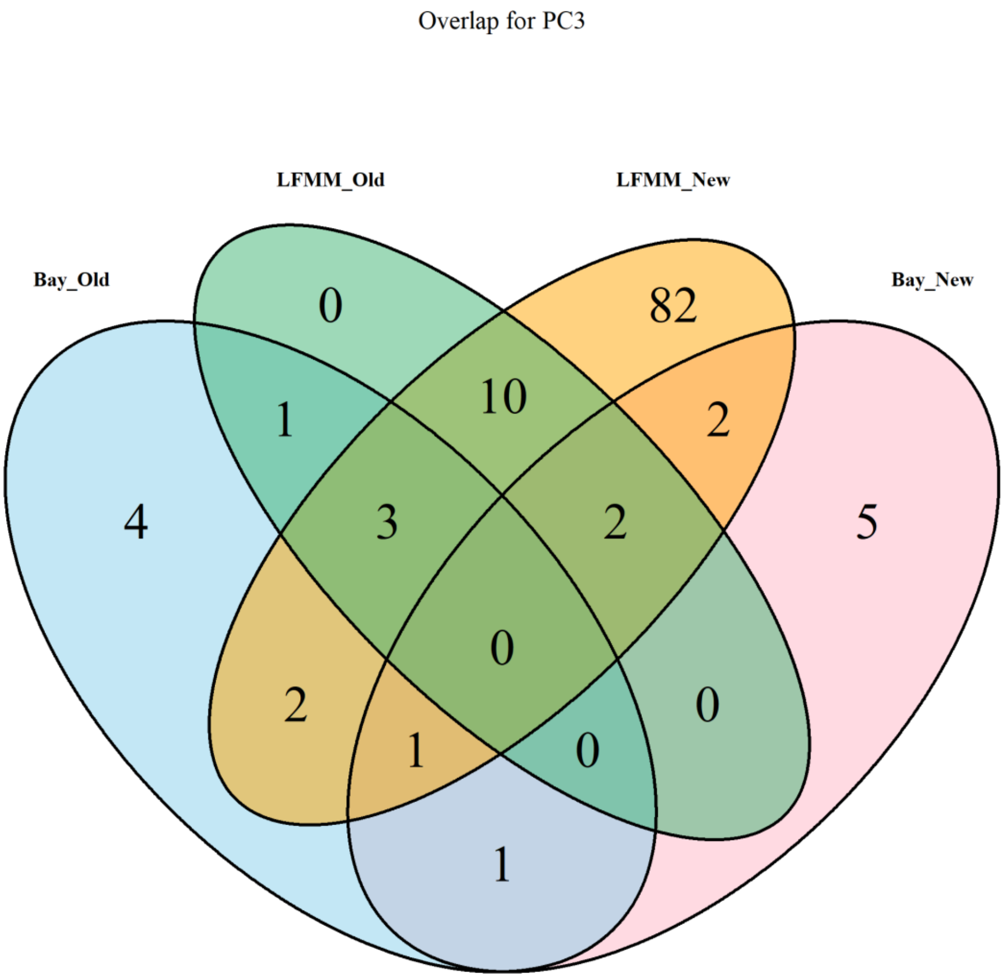


**Figure S17. Overlap of identified SNPs between the original bayenv2 and LFMM2 analyses (with MAS = 56 filter), vs. the new bayenv2 and LFMM2 analyses (with MAS = 15 filter) on PC3 calculated over all selected variables.**

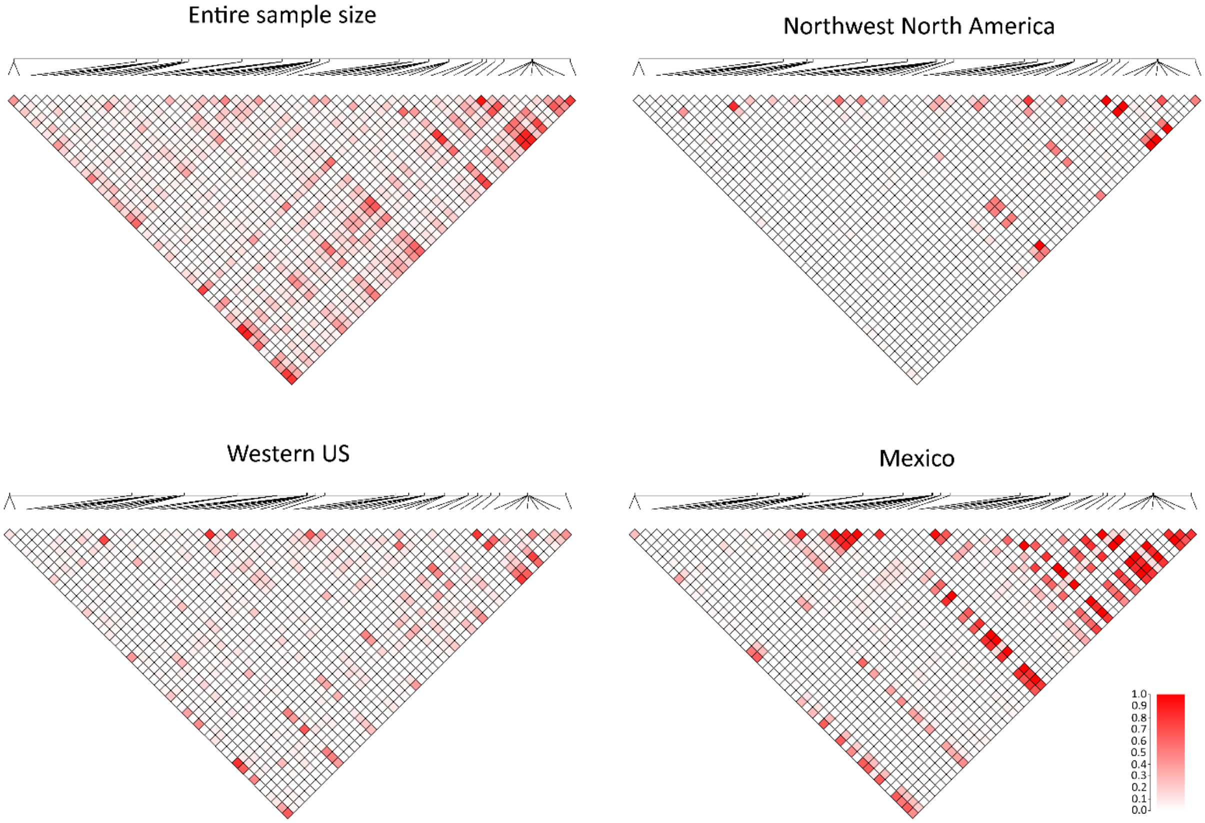


**Figure S18. Linkage-disequilibrium (LD, $r^2$) plots of SNPs identified in GEA analyses for climate PC1, PC2 and PC3 using Bayenv2 and LFMM2 (rangewide scale).** Genomic region 14611739 – 16388003 on scaffold 3 is visualized. Here, the linkage disequilibrium plots were calculated over the entire sample as well as within the northwest North America (NW-NA), western US (WU) and Mexico (MX) genetic lineages. The LD-plot for the northeast North America (NE-NA) genetic lineage, along with the annotation of coordinates, is visualized in **Fig. 5** (main text). With ranging color scheme of $r^2$=0 in white towards $r^2$=1 in red.

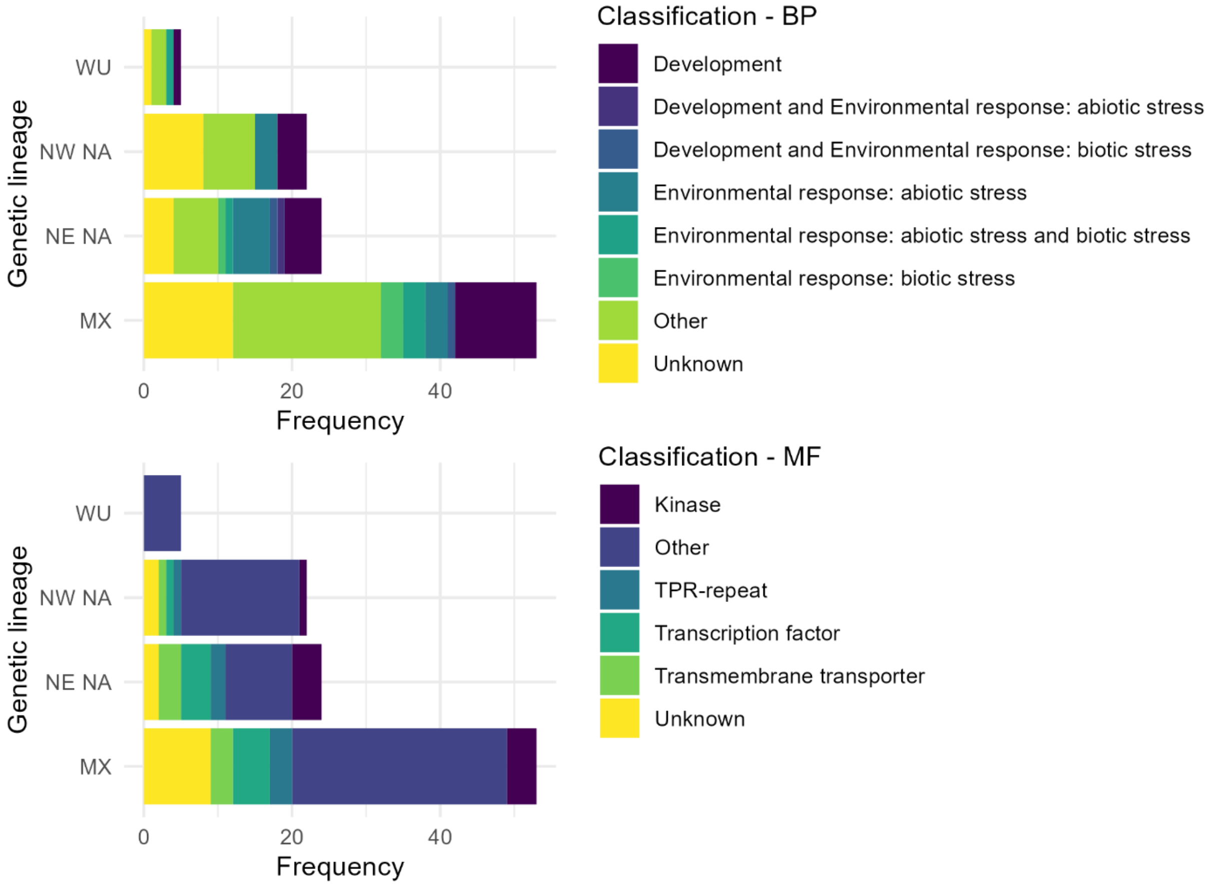


**Figure S19. Classification of genes containing the SNPs or being near the SNPs that were associated with environmental PC1, PC2 and PC3, identified within each genetic lineage.** Classification was based on literature research, separately for biological process (BP) and molecular function (MF). Genetic lineages include western US (WU), Mexico (MX), northwest North America (NW-NA) and northeast North America (NE-NA).

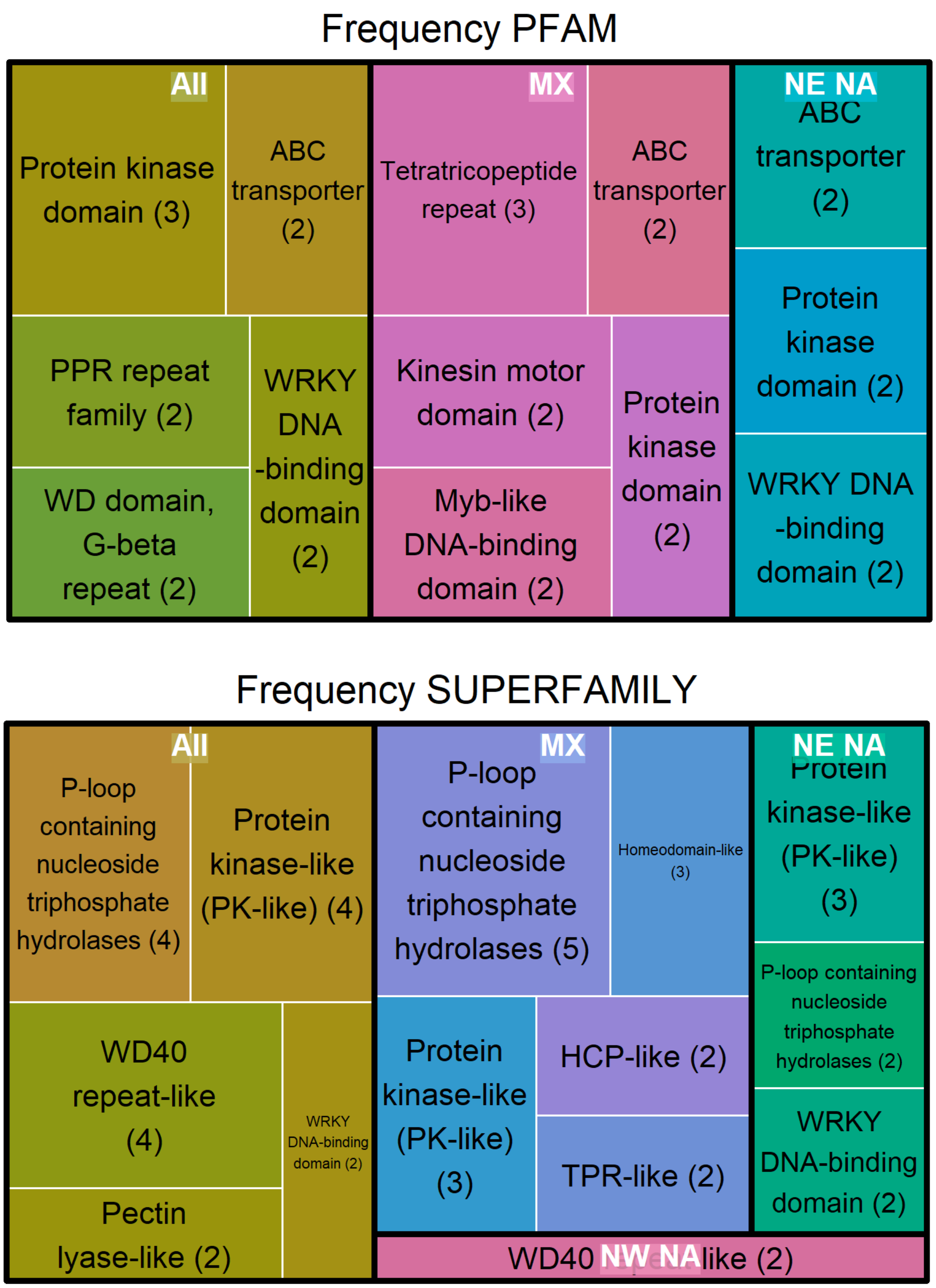


**Figure S20. Frequency of PFAMs and SUPERFAMILYs for genes with or near SNPs identified for PC1, PC2 and PC3 within each genetic lineage or over the entire sample size.** Only occurrences of at least 2 are plotted. Genetic lineages include Mexico (MX), northeast North America (NE-NA) and northwest North America (NW-NA). The western US lineage did not contain any categories occurring more than once and is thus not visualized. Actual frequencies are displayed in brackets.

**Table S1. Comparison of SNP calling with old and new reference genome without any filtering.** Total number of SNPs indicates the number of SNPs identified over all samples, which is separated for the new genome in scaffolds, contigs and haplotigs. Missingness indicates the proportion of missing data per individual (higher percentage means more missing data). Moreover, the missingness per individual for SNPs that are not mapped to haplotigs was calculated and indicated in brackets. MX indicates the Mexico genetic lineage, WU indicates the western US genetic lineage, NW-NA indicates the northwest North America lineage and NE-NA indicates the northeast North America lineage.

| | **New genome** | **Old genome** |
|---|---|---|
| Average mapped reads per sample | 3,689,432 | 3,725,816 |
| Average unmapped reads per sample | 396,460 | 499,149 |
| Average number of loci per sample | 44,175 | 42,911 |
| **Total number SNPs (raw)** | 1,666,993 | 1,556,064 |
| Scaffolds | 1,460,666 | - |
| Contigs | 22,292 | - |
| Haplotigs | 184,035 | - |
| **Average missingness per individual for unfiltered SNPs, with and [without haplotigs]** | 0.457 [0.446] | 0.4345 |
| Standard deviation over all individuals | 0.122 [0.123] | 0.1221 |
| Lineage MX | 0.461 [0.450] | 0.4474 |
| Lineage WU | 0.486 [0.477] | 0.4628 |
| Lineage NW-NA | 0.444 [0.433] | 0.4153 |
| Lineage NE-NA | 0.443 [0.431] | 0.4099 |

**Table S2. Filtering settings for dataset Set_01 (used for $F_{ST}$, phylogeny and historical gene flow analyses) and Set_02 (used for genotype-environment association (GEA) analyses).** MAS indicates minimum number of samples with rare allele. MX indicates the Mexico genetic lineage, WU indicates the western US genetic lineage, NW-NA indicates the northwest North America lineage and NE-NA indicates the northeast North America lineage.

| | **Set_01** | **Set_02** |
|---|---|---|
| **Nb individuals raw** | 1,246 | 1,689 |
| **Nb SNPs raw** | 1,666,993 | 1,666,993 |
| **Libraries** | Lib1 | Lib1+Lib2 |
| **Diploids only** | yes | yes |
| **Minimum depth** | 8 | 8 |
| **Missingness** | 80 | 90 |
| **MAS** | 31 (2.5% of samples) | 56 (3.3% of all samples) |
| **Relatedness** | 0.3 | 0.3 |
| **Linkage pruning** | r2 of 0.1 over 50 kb window | |
| **Final number SNPs** | 20,164 | 13,360 |
| **Final number individuals** | 627 | 908 |
| **Lineage MX** | 131 | 223 |
| **Lineage WU** | 67 | 132 |
| **Lineage NW-NA** | 156 | 198 |
| **Lineage NE-NA** | 273 | 355 |
| **Purpose** | Demographic history | GEA |

**Table S3. Abbreviations and explanations of environmental variables of ClimateNA database.**

| Variable | Explanation |
|---|---|
| MAT | Mean annual temperature (°C) |
| MWMT | Mean warmest month temperature (°C) |
| MCMT | Mean coldest month temperature (°C) |
| TD | Temperature difference between MWMT and MCMT (°C) |
| MAP | Mean annual precipitation (mm) |
| MSP | May to September precipitation (mm) |
| AHM | Annual heat-moisture index (MAT+10)/(MAP/1000) |
| SHM | Summer heat-moisture index (MWMT/(MSP/1000) |
| DD_0 | Degree-days below 0°C |
| DD5 | Degree-days above 5°C |
| DD_18 | Degree-days below 18°C |
| DD18 | Degree-days above 18°C |
| NFFD | Number of frost-free days |
| FFP | Frost-free period |
| bFFP | The day of the year on which FFP begins |
| eFFP | The day of the year on which FFP ends |
| PAS | Precipitation as snow (mm) |
| EMT | Extreme minimum temperature over 30 years (°C) |
| EXT | Extreme maximum temperature over 30 years (°C) |
| Eref | Hargreaves reference evaporation (mm) |
| CMD | Hargreaves climatic moisture deficit (mm) |
| RH | Mean annual relative humidity (%) |
| CMI | Hogg's climate moisture index (mm) |
| DD1040 | Degree-days above 10°C and below 40°C |

**Table S4. Statistical results for best model runs of demographic scenarios tested within single lineages with fastsimcoal2 using "bounded" keyword for upper range in time parameter (max 2000 years).** Bold rows indicate the most likely demographic scenario for each lineage based on lowest AIC (Akaike Information Criteria) values. DeltaL indicates the difference between the maximum observed likelihood (under a perfect fit of the expected to the observed SFS) and maximum estimated likelihood under the model parameters. Analysed genetic lineages included northeast North America (NE-NA), northwest North America (NW-NA), western US (WU) and Mexico (MX). Np indicates number of parameters included in the model. NCUR indicates the current population size, TG indicates the number of generations since start of expansion/contraction, NANC indicates ancient population size, GROWTHRT indicates the growth rate, NBOT indicates population size during the bottleneck, TBOT indicates the number of generations at which the bottleneck started. TG_15 indicates the generation time of TG in number of years (TG * 15 years). TBOT _15 indicates the generation time of TBOT in number of years (TBOT * 15 years).

| Lineage | Scenario | Np | deltaL | AIC | NCUR | NANC | TG | TG_15 | GROWTHRT | NBOT | TBOT | TBOT_15 |
|---|---|---|---|---|---|---|---|---|---|---|---|---|
| NE-NA | Stable | 1 | 3888.3 | 213619.0 | 108427.0 | | | | | | | |
| | Bottleneck | 4 | 997.2 | 200304.9 | 340107.0 | 1195730.0 | | | | 1423.0 | 12.0 | 180.0 |
| | Contraction | 4 | 988.4 | 200264.8 | 12292.0 | 144416.0 | 1009.0 | 15135.0 | 0.00244 | | | |
| | **Expansion** | **4** | **878.9** | **199760.5** | **14115.0** | **10146.0** | **1946.0** | **29190.0** | **-0.00017** | | | |
| NW-NA | Stable | 1 | 2735.2 | 184628.0 | 164495.0 | | | | | | | |
| | Bottleneck | 4 | 718.0 | 175338.5 | 262779.0 | 1261888.0 | | | | 1538.0 | 14.0 | 210.0 |
| | Contraction | 4 | 712.7 | 175314.0 | 16071.0 | 411819.0 | 1617.0 | 24255.0 | 0.00201 | | | |
| | **Expansion** | **4** | **604.3** | **174814.7** | **14139.0** | **10162.0** | **1955.0** | **29325.0** | **-0.00017** | | | |
| WU | Stable | 1 | 393.7 | 127767.4 | 222579.0 | | | | | | | |
| | Bottleneck | 4 | 67.0 | 126262.8 | 118205.0 | 596362.0 | | | | 3491.0 | 13.0 | 195.0 |
| | Contraction | 4 | 66.0 | 126258.5 | 18438.0 | 358275.0 | 1005.0 | 15075.0 | 0.00295 | | | |
| | **Expansion** | **4** | **25.0** | **126069.4** | **15322.0** | **10132.0** | **1808.0** | **27120.0** | **-0.00023** | | | |
| MX | Stable | 1 | 490.9 | 101408.9 | 90270.0 | | | | | | | |
| | **Bottleneck** | **4** | **9.2** | **99190.8** | **97597.0** | **227128.0** | | | | **1228.0** | **1187.0** | **17805.0** |
| | Contraction | 4 | 92.4 | 99573.9 | 63340.0 | 102709.0 | 1400.0 | 21000.0 | 0.00035 | | | |
| | Expansion | 4 | 91.4 | 99569.3 | 58544.0 | 53686.0 | 694.0 | 10410.0 | -0.00012 | | | |

**Table S5. Scenario prediction votes of Run1 using a forest of 1500 trees, in DIY-ABC-RF.** Scenario #2 is the most likely scenario as it holds the most votes, *i.e.* 709. The posterior probability of the most likely scenario was 0.62.

| Scenario | 1 | 2 | 3 | 4 | 5 | 6 | 7 | 8 | 9 | 10 |
|---|---|---|---|---|---|---|---|---|---|---|
| Votes | 654 | **709** | 23 | 9 | 30 | 19 | 27 | 13 | 6 | 10 |

**Table S6. Scenario prediction votes of Run2 using a forest of 1500 trees, in DIY-ABC-RF.** Scenario #2 is the most likely scenario as it holds the most votes, *i.e.* 370. The posterior probability of the most likely scenario was 0.625.

| Scenario | 1 | 2 | 3 | 4 | 5 | 6 | 7 | 8 | 9 | 10 | 11 | 12 | 13 |
|---|---|---|---|---|---|---|---|---|---|---|---|---|---|
| Votes | 266 | **370** | 54 | 73 | 149 | 110 | 19 | 31 | 5 | 22 | 32 | 10 | 359 |

**Table S7. The contingency table of true vs. predicted scenarios for each sample in the training set for Run1, generated by DIY-ABC-RF.** The mean classification error over all scenarios was 0.22.

| | Scenario | 1 | 2 | 3 | 4 | 5 | 6 | 7 | 8 | 9 | 10 | Class error |
|---|---|---|---|---|---|---|---|---|---|---|---|---|
| predicted | 1 | 1284 | 651 | 0 | 1 | 20 | 0 | 13 | 0 | 0 | 1 | 0.348223 |
| predicted | 2 | 650 | 1289 | 1 | 2 | 27 | 0 | 41 | 0 | 0 | 0 | 0.358706 |
| predicted | 3 | 1 | 2 | 2012 | 3 | 0 | 0 | 0 | 0 | 0 | 1 | 0.003467 |
| predicted | 4 | 0 | 1 | 8 | 1968 | 0 | 0 | 0 | 0 | 1 | 0 | 0.005056 |
| predicted | 5 | 29 | 24 | 0 | 0 | 1232 | 6 | 646 | 5 | 0 | 2 | 0.366255 |
| predicted | 6 | 0 | 0 | 0 | 0 | 8 | 1303 | 6 | 581 | 26 | 22 | 0.330421 |
| predicted | 7 | 20 | 27 | 0 | 0 | 708 | 8 | 1319 | 11 | 0 | 0 | 0.369804 |
| predicted | 8 | 0 | 0 | 0 | 0 | 4 | 629 | 15 | 1365 | 26 | 16 | 0.335766 |
| predicted | 9 | 0 | 0 | 0 | 0 | 0 | 19 | 0 | 30 | 1909 | 33 | 0.041185 |
| predicted | 10 | 0 | 0 | 1 | 0 | 0 | 32 | 0 | 25 | 40 | 1896 | 0.049147 |

**Table S8. The contingency table of true vs. predicted scenarios for each sample in the training set for Run2, generated by DIY-ABC-RF.** The mean classification error over all scenarios was 0.48.

| | Scenario | 1 | 2 | 3 | 4 | 5 | 6 | 7 | 8 | 9 | 10 | 11 | 12 | 13 | Class error |
|---|---|---|---|---|---|---|---|---|---|---|---|---|---|---|---|
| predicted | 1 | 1292 | 665 | 34 | 54 | 58 | 19 | 3 | 4 | 0 | 4 | 146 | 131 | 228 | 0.510235 |
| predicted | 2 | 509 | 946 | 25 | 33 | 30 | 2 | 0 | 0 | 0 | 0 | 121 | 80 | 158 | 0.503151 |
| predicted | 3 | 2 | 1 | 1088 | 159 | 0 | 470 | 236 | 81 | 250 | 80 | 4 | 5 | 0 | 0.542088 |
| predicted | 4 | 8 | 1 | 142 | 1172 | 70 | 241 | 339 | 89 | 169 | 119 | 4 | 1 | 7 | 0.50381 |
| predicted | 5 | 15 | 9 | 5 | 50 | 1419 | 19 | 6 | 325 | 2 | 235 | 5 | 2 | 14 | 0.326211 |
| predicted | 6 | 0 | 0 | 373 | 82 | 0 | 923 | 249 | 59 | 224 | 101 | 1 | 1 | 0 | 0.54148 |
| predicted | 7 | 0 | 0 | 73 | 248 | 3 | 178 | 635 | 77 | 344 | 85 | 0 | 0 | 0 | 0.613512 |
| predicted | 8 | 5 | 1 | 7 | 36 | 237 | 15 | 38 | 842 | 10 | 440 | 12 | 13 | 5 | 0.493076 |
| predicted | 9 | 0 | 0 | 120 | 183 | 0 | 140 | 444 | 24 | 988 | 72 | 0 | 0 | 0 | 0.498732 |
| predicted | 10 | 0 | 0 | 0 | 32 | 108 | 33 | 59 | 489 | 23 | 830 | 1 | 0 | 5 | 0.474684 |
| predicted | 11 | 76 | 70 | 25 | 4 | 6 | 7 | 2 | 9 | 0 | 2 | 741 | 769 | 12 | 0.569936 |
| predicted | 12 | 73 | 61 | 40 | 8 | 21 | 2 | 4 | 17 | 0 | 2 | 868 | 904 | 11 | 0.550472 |
| predicted | 13 | 69 | 179 | 19 | 31 | 23 | 37 | 2 | 5 | 2 | 1 | 26 | 32 | 1586 | 0.21173 |

**Table S9. Number of identified environmentally associated SNPs per scaffold for all 24 environmental variables and PC1, PC2 and PC3, based on the entire sample size.** Scaffold indicates the scaffold number. Total indicates the number of total filtered SNPs called on this scaffold. B_SNPs indicates the number of SNPs identified by Bayenv2 on this scaffold along with the represented percentage (B_%). L_SNPs indicates the number of SNPs identified by LFMM2 on this scaffold along with the represented percentage (L_%). Top three scaffolds with highest representation of adaptive SNPs are in bold.

| Scaffold | Total | B_SNPs | B_% | L_SNPs | L_% |
|---|---|---|---|---|---|
| 20g1 | 1057 | 21 | 2 | 23 | 2.2 |
| 20g2 | 1047 | 36 | 3.4 | 38 | 3.6 |
| **20g3** | **1187** | **70** | **5.9** | **78** | **6.6** |
| 20g5 | 798 | 17 | 2.1 | 29 | 3.6 |
| 20g6 | 384 | 3 | 0.8 | 6 | 1.6 |
| 20g7 | 964 | 24 | 2.5 | 36 | 3.7 |
| 20g8 | 918 | 15 | 1.6 | 38 | 4.1 |
| 20g9 | 335 | 7 | 2.1 | 7 | 2.1 |
| 20g10 | 1037 | 24 | 2.3 | 33 | 3.2 |
| 20g11 | 273 | 6 | 2.2 | 4 | 1.5 |
| 20g12 | 620 | 12 | 1.9 | 18 | 2.9 |
| **20g13** | **1045** | **39** | **3.7** | **57** | **5.5** |
| 20g14 | 496 | 10 | 2 | 5 | 1 |
| **20g15** | **473** | **24** | **5.1** | **21** | **4.4** |
| 20g16 | 377 | 6 | 1.6 | 7 | 1.9 |
| 20g17 | 504 | 12 | 2.4 | 12 | 2.4 |
| 20g18 | 453 | 2 | 0.4 | 10 | 2.2 |
| 20g19 | 541 | 8 | 1.5 | 5 | 0.9 |
| 20g20 | 648 | 12 | 1.9 | 26 | 4 |

**Table S10. Number of overlapping SNPs between the original runs (Bayenv2 and LFMM2 with MAS=56 filter) vs. the new runs (Bayenv2 and LFMM2 with MAS=15 filter) for PC1-2-3 calculated on all environmental variables.** The first four rows indicate the total number of SNPs identified.

| Analysis | PC1 | PC2 | PC3 |
|---|---|---|---|
| Bayenv_56 | 50 | 46 | 12 |
| Bayenv_15 | 39 | 53 | 15 |
| LFMM_56 | 18 | 6 | 16 |
| LFMM_15 | 15 | 53 | 102 |
| Bayenv_56 vs. Bayenv_15 | 23 | 26 | 2 |
| Bayenv_56 vs. LFMM_56 | 14 | 4 | 4 |
| Bayenv_56 vs. LFMM_15 | 8 | 30 | 6 |
| Bayenv_15 vs. LFMM_56 | 13 | 4 | 2 |
| Bayenv_15 vs. LFMM_15 | 8 | 30 | 5 |
| LFMM_56 vs. LFMM_15 | 11 | 4 | 15 |
| Bayenv_56 vs. Bayenv_15 vs. LFMM_56 | 13 | 4 | 0 |
| Bayenv_56 vs. Bayenv_15 vs. LFMM_15 | 8 | 25 | 1 |
| Bayenv_56 vs. LFMM_56 vs. LFMM_15 | 8 | 4 | 3 |
| Bayenv_15 vs. LFMM_56 vs. LFMM_15 | 8 | 4 | 2 |
| Overlap all four | 8 | 4 | 0 |

**Table S11. Number of SNPs and genes identified with LFMM2 and Bayenv2 per tested climatic variable within each genetic lineage.** ENV indicates the environmental variables, principal components (PC), as well as the library variable (LIB, only tested for with LFMM2). Lineage indicates the genetic lineage. SNPs_B indicates the total number of SNPs identified using Bayenv2. SNPs_L B indicates the total number of SNPs identified using LFMM2. SNPs_O indicates the overlap between SNPs_B and SNPs_L. Genes_B indicates the number of unique genes that SNPs were identified within 5kb proximity using Bayenv2. Genes_B indicates the number of unique genes that SNPs were identified within 5kb proximity using LFMM2. Genes_O indicates the overlap between Genes_B and Genes_L. UN_B indicates the number of SNPs that are unique to the specified climate variable in respect to all SNPs identified by Bayenv2. UN_L indicates the number of SNPs that are unique to the specified climate variable in respect to all SNPs identified by LFMM2. Numbers in brackets for the NW-NA lineage are those identified with only the first GBS library using LFMM2. MX indicates the Mexico genetic lineage, WU indicates the western US genetic lineage, NW-NA indicates the northwest North America lineage and NE-NA indicates the northeast North America lineage.

| ENV | Lineage | SNPs_B | SNPs_L | SNPs_O | Genes_B | Genes_L | Genes_O | Un_B | Un_L |
|---|---|---|---|---|---|---|---|---|---|
| PC1 | NE-NA | 3 | 25 | 2 | 3 | 15 | 2 | 3 | 24 |
| PC2 | NE-NA | 1 | 2 | 0 | 1 | 2 | 0 | 1 | 1 |
| PC3 | NE-NA | 1 | 0 | 0 | 1 | 0 | 0 | 1 | 0 |
| LIB | NE-NA | | 0 | | | 0 | | | 0 |
| PC1 | NW-NA | 1 | 72 [17] | 0 | 1 | 44 [12] | 0 | 1 | 42 [14] |
| PC2 | NW-NA | 0 | 36 [4] | 0 | 0 | 23 [3] | 0 | 0 | 9 [0] |
| PC3 | NW-NA | 0 | 5 [3] | 0 | 0 | 3 [1] | 0 | 0 | 5 [4] |
| LIB | NW-NA | | 24 | | | 10 | | | 15 |
| PC1 | WU | 2 | 0 | 0 | 2 | 0 | 0 | 2 | 0 |
| PC2 | WU | 1 | 0 | 0 | 1 | 0 | 0 | 1 | 0 |
| PC3 | WU | 2 | 0 | 0 | 2 | 0 | 0 | 2 | 0 |
| LIB | WU | | 0 | | | 0 | | | 0 |
| PC1 | MX | 6 | 2 | 0 | 6 | 2 | 0 | 6 | 2 |
| PC2 | MX | 1 | 0 | 0 | 1 | 0 | 0 | 1 | 0 |
| PC3 | MX | 8 | 50 | 7 | 7 | 28 | 3 | 8 | 48 |
| LIB | MX | | 0 | | | 0 | | | 0 |

**Methods S1.** We compared SNP calling of the new vs. old reference genome (popgenie.org, v1.1), based on several aspects, including the average number of reads that were mapped across all samples, the average number of unmapped reads and the average number of called loci over all samples, the number of overall called SNPs and sample missingness. A comparison of the SNP calling between the old reference genome (Lin *et al*., 2018) and our new reference genome is shown in **Table S1**. Although there is no improvement in the average missingness per individual, the overall number of SNPs increased by more than 100k, however these were all mapped against haplotigs.

**Methods S2.** Fastsimcoal is a coalescent-based genetic simulation program (Excoffier *et al*., 2021a). As input, the program uses the SFS, a template file (.tpl) that defines the model, as well as an estimate file (.est) that defines the initial search ranges for the required parameters (such as population size and timepoints). While the lower boundary of the search range is respected, there was no upper range limit and thus the model could drift outside the initial predefined range (Excoffier *et al*., 2021b), except when one uses the "bounded" keyword in the .est file. First, we ran the models without using the "bounded" keyword. Hereafter, we ran the models a second time, using the parameter "bounded" for the tested time range, so that the upper range of the parameter was bounded during parameter estimation. When using the bounded keyword to constrain the upper time limit to 2,000 generations (**Table S4**), we found that the NE-NA, NW-NA and WU lineages experienced expansion since respectively 1,946, 1,955 and 1,808 generations, *i.e*. 29,190, 29,325 and 27,120 years ago. The 29 kyrs (thousand years) ago is at the upper limit of the search range, and therefore we think the run without the "bounded" parameter is likely most accurate. Therefore, the run without the "bounded" parameter is presented in the main document.

**Methods S3.** The program implements approximate Bayesian computing (ABC) with supervised machine learning based on random forests (RF). DIY-ABC-RF allows for complex scenarios, including population divergence, change in population size and admixture events, but does not allow for continuous gene flow. First, a training set must be created. This training set is the result of simulations of the different scenarios, available SNPs and sampled individuals under parameter values drawn from prior distributions that are set by the user. The simulations are based on coalescence theory. Each resulting set is summarized using various descriptive statistics. A statistical analysis based on the RF algorithms of the observed dataset versus the training dataset is then calculated to select the most likely scenario and parameter values. The RF scenario choice yields a classification vote for each scenario, representing the number of times a scenario is selected in a forest of n trees, and the scenario with most votes is the most compatible scenario to the observed dataset among the set of tested scenarios.

**Methods S4.** To assess the impact of a less stringent minimum allele sample size (MAS), we tested a lower threshold of 15 individuals (compared to the original MAS of 56). This adjustment resulted in a dataset comprising 923 individuals and 26,696 SNPs, of which 13,002 SNPs were also present in the original dataset (MAS = 56). For the MAS = 15 dataset, we conducted a correlation analysis to identify sets of environmental variables with pairwise correlations below 0.8. Two variables, EXT and DD1040, were retained despite slightly higher correlations (0.82 and 0.84, respectively) due to their ecological relevance. Principal components (PC1, PC2, and PC3) were calculated both from all environmental variables and from the subset of selected uncorrelated variables. These principal components were then used as environmental predictors in Bayenv2 and LFMM2 analyses.

**Methods S5.** Environmental association analysis was performed using LFMM2 and Bayenv2. LFMM2 simultaneously infers population structure using latent factors, *i.e.* unobserved variables. Bayenv2 calculates a covariance matrix to account for differences that represent neutral population structure and is included as a null model in the subsequent step of Bayenv2. The relatively low number of overlapping SNPs per climate variable as identified between LFMM2 and Bayenv2 analyses, over the entire sample size as well as on a lineage basis, could be due to numerous factors. Both methods use a form of mixed effect models that provide control of neutral population structure, in which the allelic frequencies are response variables, environmental data are fixed factors and neutral population structure is treated as a random factor, and both use a linear type of genotype-environment association (Rellstab *et al*., 2015). However, the methods differ in significance testing, and in attesting for neutral population structure, *i.e.* Bayenv2 uses a covariance matrix of estimated allele frequencies, while LFMM2 uses random factors (latent factors) which are similar to principal component and calculated over all SNPs (Rellstab *et al*., 2015). Additionally, LFMM2 does not allow for missing data, and thus, only works with imputed data, which potentially increases the number of false positives (Caye *et al.*, 2019). LFMM2 was calculated on an individual basis, while Bayenv2 bases calculation on a population basis while correcting for population size. Moreover, LFMM was found to better detect signatures of weak selection in comparison to Bayenv (Frichot *et al.*, 2013). Lastly, LFMM2 was run within each genetic lineage using all 24 environmental variables (**Dataset S9**).

**Results S1.** Based on correlation analyses among environmental variables (**Fig. S9**), we retained the following for downstream analyses: MWMT, MCMT, MAP, MSP, FFP, AHM, PAS, EXT, RH, and DD1040. These represent: Mean Warmest Month Temperature (°C), Mean Coldest Month Temperature (°C), Mean Annual Precipitation (mm), May–September Precipitation (mm), Frost-Free Period, Annual Heat-Moisture Index ((MAT + 10)/(MAP/1000)), Precipitation as Snow (mm), Extreme Maximum Temperature (°C), Mean Annual Relative Humidity (%), and Degree-Days above 10°C and below 40°C, respectively (**Fig. S10**).

The distribution of environmental variables along the first three principal components closely mirrored that of the original dataset (**Figs. S11a-S12a** compared to **Fig. S7a**). When principal components were computed using only the selected uncorrelated variables, the sample distribution pattern remained largely consistent (**Figs. S11b-S12b** compared to **Fig. S8a**). Notably, individuals from the MX lineage exhibited the most distinct environmental profiles compared to other genetic clusters. Identified significant SNP distribution patterns were also highly comparable between the datasets filtered with MAS = 56 and MAS = 15, with both showing prominent clustering of SNPs on scaffolds 3 and 13 (**Fig. S13**). Similar congruence was observed when assessing the principal components calculated on the selected uncorrelated variables (**Fig. S14**). Given the similarity across approaches, we proceeded with analyses using principal components derived from the full set of environmental variables. The number of SNPs identified by Bayenv2 remained relatively stable between MAS thresholds: 50, 46, and 12 SNPs for PC1, PC2, and PC3 respectively in the MAS = 56 dataset, and 39, 53, and 15 SNPs for MAS = 15. LFMM2, however, identified substantially more SNPs using the lower MAS filter (15, 53, and 102 SNPs for PC1, PC2, and PC3 respectively), compared to 18, 6, and 16 for the original filter. There was considerable overlap in SNPs identified across MAS thresholds, *i.e.* 23, 26, and 2 for respectively PC1. PC2 and PC3 for between MAS=15 and MAS=56 for bayenv2, and respectively 11, 4, and 15 between both MAF filters for LFMM2 (see **Table S10** and **Figs. S15–S17**).

The following datasets and files will be uploaded as Excel spreadsheets:

**Dataset S1.** Overview of sample collection in this study. This includes individual names, population names, geographical coordinates, species, library number, and ploidy assignment.

**Dataset S2.** Overview of climate variables used in the genotype-environment associations.

**Dataset S3.** SNPs identified for all environmental variables using Bayenv2 for entire sample size. Explanation of column names are provided in the spreadsheet.

**Dataset S4.** SNPs identified for all environmental variables using LFMM2 for entire sample size. Explanation of column names are given in the spreadsheet.

**Dataset S5.** SNPs identified for PC1, PC2 and PC3 using Bayenv2 and LFMM2 along the entire sample size. Explanation of column names are provided within the spreadsheet.

**Dataset S6.** Inversion data of four genetically distinct individuals.

**Dataset S7.** SNPs identified for PC1, PC2 and PC3 using Bayenv2 and LFMM2 within each respective genetic lineage. Explanation of column names are found in the spreadsheet.

**Dataset S8.** Annotation summary of SNPs identified for PC1, PC2 and PC3 using Bayenv2 and LFMM2 within each respective lineage.

**Dataset S9.** SNPs identified for all environmental variables using LFMM2 within each respective lineage. Explanation of column names are given within the spreadsheet.

**headerRF_run1.txt** The input file for DIY-ABC-RF2 for run1. The file described all tested scenarios and input parameters.

**headerRF_run2.txt** The input file for DIY-ABC-RF2 for run2. The file described all tested scenarios and input parameters.
